# TIN2 facilitates TRF1-mediated *trans*- and *cis*-interactions on physiologically relevant long telomeric DNA

**DOI:** 10.1101/2020.09.12.286559

**Authors:** Hai Pan, Parminder Kaur, Ming Liu, Pengning Xu, Chelsea Mahn, Ryan Barnes, Qingyu Tang, Pengyu Hao, Dhruv Bhattaram, Changjiang You, Jacob Piehler, Keith Weninger, Robert Riehn, Susan Smith, Patricia L. Opresko, Hong Wang

**Affiliations:** Physics Department, North Carolina State University, Raleigh, North Carolina, 27695, USA; Center for Human Health and the Environment, North Carolina State University, Raleigh, North Carolina, 27695, USA; Department of Environmental and Occupational Health, UPMC Hillman Cancer Center, University of Pittsburgh, Pittsburgh, Pennsylvania 15219, USA; Department of Molecular and Structural Biochemistry, North Carolina State University, Raleigh, North Carolina, 27695, USA; Department of Biomedical Engineering, Georgia Institute of Technology & Emory University of Medicine, Atlanta, Georgia, 30332, USA; Division of Biophysics, Universität Osnabrück, Barbarstrasse 11, 49076, Osnabrück, Germany; Kimmel Center for Biology and Medicine at the Skirball Institute, Department of Pathology, New York University School of Medicine, New York, NY 10016, USA; Molecular Biophysics and Structural Biology Graduate Program, Carnegie Mellon University and the University of Pittsburgh, Pittsburgh, Pennsylvania, 15219, USA; Toxicology and Environmental Science Program, North Carolina State University, Raleigh, North Carolina, 27695, USA

**Keywords:** shelterin proteins, TRF1 and TIN2, protein-DNA interactions, single-molecule imaging, atomic force microscopy

## Abstract

The shelterin complex consisting of TRF1, TRF2, RAP1, TIN2, TPP1, and POT1, functions to prevent false recognition of telomeres as double-strand DNA breaks, and to regulate telomerase and DNA repair protein access. TIN2 is a core component linking double-stranded telomeric DNA binding proteins (TRF1 and TRF2) and proteins at the 3’ overhang (TPP1-POT1). Since knockdown of TIN2 also removes TRF1 and TRF2 from telomeres, determining TIN2’s unique mechanistic function has been elusive. Here, we investigated DNA molecular structures promoted by TRF1-TIN2 using complementary single-molecule imaging platforms, including atomic force microscopy (AFM), total internal reflection fluorescence microscopy (TIRFM), and the DNA tightrope assay. We demonstrate that TIN2S and TIN2L isoforms facilitate TRF1-mediated DNA compaction (*cis*-interactions) and DNA-DNA bridging (*trans*-interactions) in a telomeric sequence- and length-dependent manner. On the short telomeric DNA substrate (6 TTAGGG repeats), the majority of TRF1 mediated telomeric DNA-DNA bridging events are transient with a lifetime of ~1.95 s. On longer DNA substrates (270 TTAGGG), TIN2 forms multi-protein complexes with TRF1 and stabilizes TRF1-mediated DNA-DNA bridging events that last for at least minutes. Preincubation of TRF1 with its regulator protein Tankyrase 1 significantly reduces TRF1-TIN2 mediated DNA-DNA bridging, whereas TIN2 protects the disassembly of TRF1-TIN2 mediated DNA-DNA bridging upon Tankyrase 1 addition. Our study provides evidence that TIN2 functions to promote TRF1 mediated *trans*-interactions of telomeric DNA, leading to new mechanistic insight into sister telomere cohesion.

## INTRODUCTION

Telomeres are nucleoprotein structures that prevent the degradation or fusion of the ends of linear chromosomes, which are threatened by at least seven distinct DNA damage response (DDR) pathways (1–3). Human telomeres contain ~2 to 20 kb of TTAGGG repeats and a G-rich 3’ overhang of ~50-400 nt in length (1,4). In humans, a specialized six-protein shelterin complex consisting of TRF1, TRF2, RAP1, TIN2, TPP1, and POT1, binds specifically to the unique sequence and structure at telomeres to protect chromosome ends. Prevention of telomeres from being falsely recognized as double-strand DNA breaks and regulation of DNA repair protein access depend on the biochemical activities of shelterin proteins and their collaborative actions with other proteins involved in the genome maintenance pathways (5–9). Extensive telomere shortening or dramatic telomere loss due to DNA damage causes de-protection, which triggers cell senescence and aging-related pathologies (10,11).

The main protein-protein and protein-DNA interactions at telomeres have been investigated using crystallography, biochemical assays, yeast two-hybrid systems, co-immunoprecipitation, as well as visualization of shelterin subcomponents *in vitro* and *in vivo* using fluorescence imaging (3). Among shelterin components, both TRF1 and TRF2 specifically recognize double-stranded telomeric DNA through the Myb/SANT domain facilitated by homodimerization through the TRFH domain (12,13). However, TRF1 and TRF2 display distinct DNA binding properties and functions. TRF1 and TRF2 contain an acidic or a basic domain, respectively, at their N-termini (14). TRF2 prevents Mre11/Rad50/Nbs1-dependent ATM kinase signaling, classical nonhomologous end-joining (NHEJ), as well as alternative nonhomologous end-joining (alt-NHEJ) pathways at telomeres. These distinct functions of TRF2 are believed to be mediated through its activities in promoting and stabilizing T-loops, in which the 3’ single-strand overhang invades the upstream double-stranded telomeric region (15–18). The TRFH domain of TRF2 binds nonspecifically to dsDNA with low affinity and mediates the wrapping of dsDNA, which is proposed to be an essential step in the T-loop formation (17). In comparison, TRF1 represses telomere fragility by preventing DNA replication fork stalling at telomeres (19). TRF1 promotes parallel pairing of telomeric DNA tracts (20,21). A flexible domain in TRF1 enables the two Myb domains in the TRF1 dimer to interact with DNA independently and to mediate looping of telomeric DNA (22).

TIN2 itself does not have binding affinity to either double-stranded or single-stranded DNA (23). However, it is a core shelterin component that bridges double-stranded (TRF1 and TRF2) and single-stranded telomeric DNA binding proteins (TPP1-POT1) (24–28). Crystal structures revealed the interfaces between TRF1-TIN2, TRF2-TIN2, and TPP1-TIN2 (29,30). TIN2 stabilizes both TRF1 and TRF2 at telomeres (31,32). The loss of the TRF1 or TRF2 binding domains in TIN2 triggers a DNA damage response (33). Binding of TPP1 to TIN2 is required for POT1-mediated telomere protection (34). As an integral component of the “TIN2/TPP1/POT1 processivity complex”, TIN2 functions together with TPP1/POT1 to stimulate telomerase processivity (35). Collectively, these results demonstrated that distinct domains of TIN2 bind to TRF1, TRF2, or TPP1 independently. Lacking either domain leads to telomere dysfunction. Furthermore, TIN2 directly interacts with the cohesin subunit SA1 and plays a key role in a distinct SA1-TRF1-TIN2 mediated sister telomere cohesion pathway that is largely independent of the cohesin ring subunits (8,36). Binding of TRF1-TIN2 to telomeres is regulated by the poly(ADP-ribose) polymerase Tankyrase 1 (37). ADP-ribosylation of TRF1 by Tankyrase 1 reduces its binding to telomeric DNA *in vitro,* and the depletion of Tankyrase 1 using siRNA leads to mitotic arrest and persistent telomere cohesion that can be rescued by depletion of TIN2 (8,36,38).

Three distinct TIN2 isoforms have been identified in human cell lines (35,39,40) that include TIN2S (354 AAs), TIN2L (451 AAs), and TIN2M (TIN2 medium, 420 AAs), with the latter retaining the last intron between exons 8 and 9. TIN2S, TIN2L, and TIN2M share the same TRF1, TRF2, and TPP1 binding domains and localize to telomeres (23,35,39). In addition, TIN2L associates strongly with the nuclear matrix (39). A recent study showed that TIN2L, but not TIN2S, is phosphorylated by the casein kinase 2 (CK2) (41). TIN2L phosphorylation enhances its association with TRF2 *in vivo,* while TRF1 interacts more robustly with TIN2S than TIN2L in the cellular environment.

Consistent with its key role in telomere maintenance, germline inactivation of TIN2 in mice is embryonic lethal (42). Removal of TIN2 leads to the formation of telomere dysfunction-induced foci (TIFs). Importantly, clinical studies further highlight the biological significance of TIN2 in telomere protection (43,44). *TINF2,* which encodes TIN2, is the second most frequently mutated gene in the telomere elongation and protection disorder dyskeratosis congenita (DKC). DKC-associated TIN2 mutations are most frequently *de novo* and cluster at a highly conserved region near the end of its TRF1-binding domain.

Two decades of research since the first discovery of TIN2 have shed light on protein-interaction networks around TIN2 and its multi-faceted roles in telomere maintenance. However, since TIN2 itself does not directly bind to DNA and instead serves as a “mediator/enhancer” for shelterin and telomerase activities, defining TIN2’s distinct function at the molecular level has been challenging. The bottle-neck for studying TIN2 lies in the fact that results from bulk biochemical assays do not fully reveal the heterogeneity and dynamics of the protein-protein and protein-DNA interactions. Furthermore, cell-based assays only provide information on the outcomes from downstream effectors after the knocking down of TIN2 that also removes TRF1 and TRF2 from telomeres. These approaches do not allow us to investigate the molecular structures and dynamics in which TIN2 directly participates. To address this long-standing question, we applied complementary single-molecule imaging platforms, including atomic force microscopy (AFM) (45–47), total internal reflection fluorescence microscopy (TIRFM) (48), and the DNA tightrope assay to monitor TRF1-TIN2 mediated DNA compaction and DNA-DNA bridging (49–51). Through using DNA substrates on different length scales (6 and 270 TAAGGG repeats), these imaging platforms provide complementary results demonstrating that both TIN2S and TIN2L facilitate TRF1-mediated DNA compaction (*cis*-interactions) and DNA-DNA bridging (*trans*-interactions) in a telomeric sequence- and length-dependent manner. On the short telomeric DNA substrate (6 TTAGGG repeats), the majority of TRF1 mediated telomeric DNA-DNA bridging events are transient with a lifetime of ~1.95 s. TIN2 stabilizes TRF1-mediated DNA-DNA bridging events that last for minutes. In some cases, TRF1-TIN2 is capable of mediating the bridging of multiple copies of telomeric DNA fragments. Importantly, our results demonstrate that TIN2 protects the disassembly of TRF1-TIN2-mediated DNA-DNA bridging by Tankyrase 1.

In summary, this study uncovered the unique biophysical function of TIN2 as a telomeric architectural protein, acting together with TRF1 to mediate interactions between distant telomeric sequences. Furthermore, this work establishes a unique combination of single-molecule imaging platforms for future examination of TIN2 disease variants and provides a new direction for investigating molecular mechanisms underlying diverse TIN2 functions.

## RESULTS

### TRF1-TIN2 promotes telomeric DNA compaction and DNA-DNA bridging

Previously, TIN2 was shown to modulate the bridging of telomeric DNA by TRF1 (31). However, the bulk biochemical assays using short telomeric DNA (six telomeric repeats) did not provide information regarding the structure of the TRF1-TIN2-DNA complex. To evaluate how TIN2 regulates TRF1 DNA binding, we applied AFM imaging to investigate how TIN2 affects the telomeric DNA-DNA pairing mediated by TRF1 at the single-molecule level on longer telomeric DNA substrates (270 TTAGGG repeats). We purified TRF1 (**Figure S1A**), and obtained TIN2S (1-354 amino acids, 39.4 kDa), and TIN2L (1-451 amino acids, 50.0 kDa) proteins purified from insect cells (**Figure 1A** and **Figure S1B**). Previously, we established an AFM imaging-based calibration method to investigate the oligomeric states and protein-protein interactions by correlating AFM volumes of proteins and their molecular weights (45,47,52). AFM volumes of TRF1 alone in solution showed two distinct peaks centered at 45.5 nm^3^ (± 12.3 nm^3^) and 105.4 nm^3^ (± 22.2 nm^3^), which were consistent with TRF1 monomers (51 KDa) and dimers (102 KDa), respectively (**Figure S1A**). Meanwhile, AFM volumes of purified TIN2S (41.3 nm^3^ ± 28.3 nm^3^) and TIN2L (41.9 ± 12.8 nm^3^) were consistent with the notion that TIN2 does not interact with itself (23), and TIN2 exists in a monomeric state in solution (**Figure S1B**). Next, to validate the activities of TIN2, we used electrophoresis mobility shift assays (EMSAs) to verify the interaction of TIN2 with TRF1 on a double-stranded telomeric DNA substrate (48 bp containing 3 TTAGGG repeats, **Figure S2**). Consistent with previous studies (23), EMSA experiments showed that TIN2S and TIN2L did not directly bind to telomeric dsDNA (**Figures S2A**). Both TRF1-TIN2S and TRF1-TIN2L induced a clear supershift of the telomeric DNA substrate compared to TRF1 alone (Complex III in **Figure S2B&C**), indicating the formation of stable TRF1-TIN2-telomeric DNA complexes.

**Figure 1.**
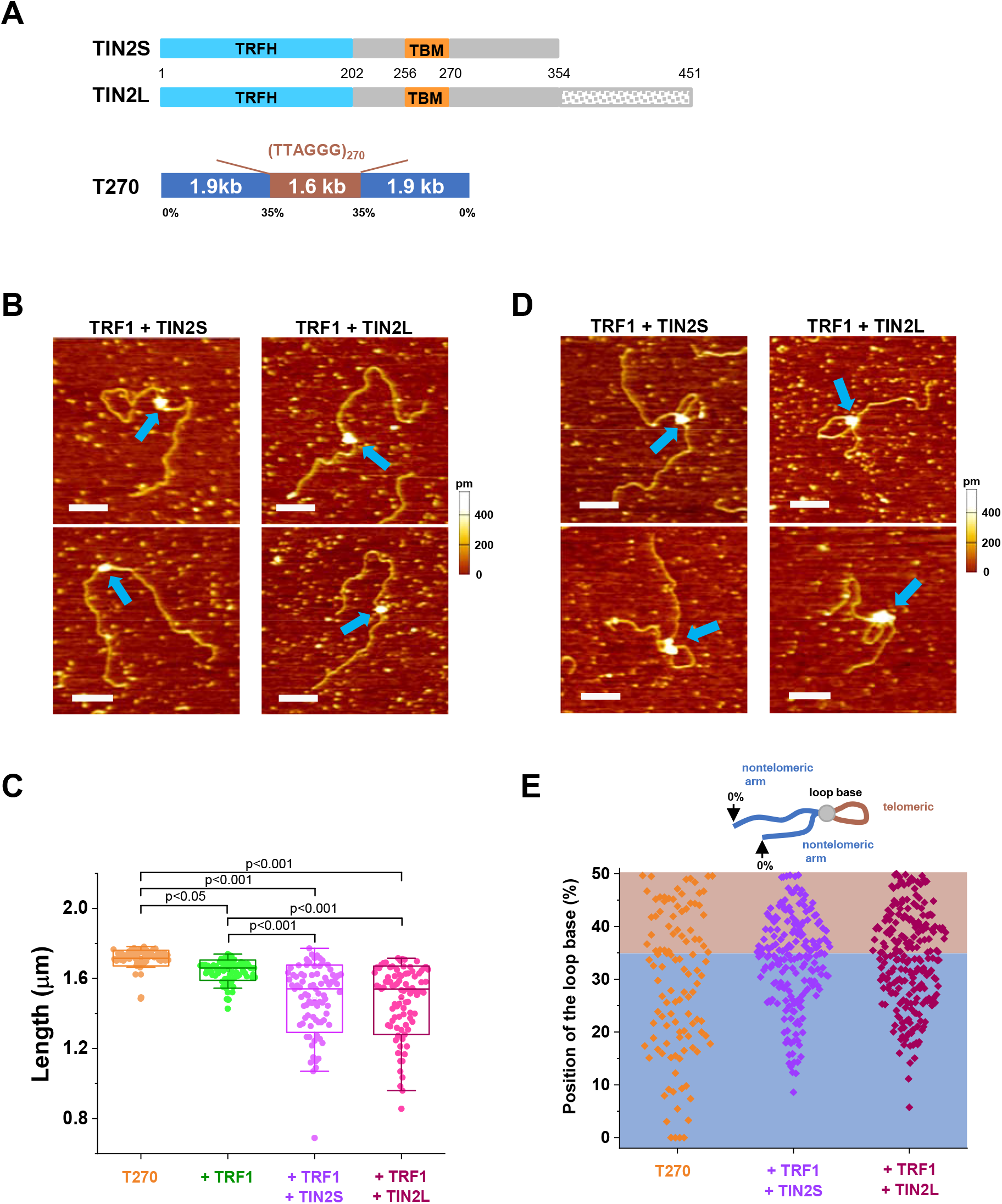
TRF1-TIN2 compacts DNA and induces DNA looping in a telomeric DNA sequence-dependent manner. (**A**) Schematics of domain structures of TIN2S and TIN2L (top panels) and the linear T270 DNA substrate (bottom panel). TRFH: telomeric repeat factor homology domain; TBM: TRF1, TRF2 binding domain. TIN2L has an additional 97 amino acids at its C-terminus. T270 DNA (5.4 kb) contains 1.6 kb (270 TTAGGG) telomeric repeats in the middle that is 35%-50% from DNA ends. (**B**) Representative AFM images showing TRF1-TIN2S (arrows, left panels) and TRF1-TIN2L (arrows, right panels) complexes on the linear T270 DNA. (**C**) TRF1-TIN2 compacted the telomeric DNA. DNA length measurements for the linear T270 DNA alone (1.71 ± 0.05 μm, N=90), T270 in the presence of TRF1 (1.64 ± 0.06 μm, N=91), TRF1 and TIN2S (1.58 ± 0.11 μm, N=85), or TRF1 and TIN2L (1.59 ± 0.1 μm, N=89). Boxes represent ±SD. Error bars indicate ranges within 1.5QR. (**D**) Representative AFM images showing DNA loop formation upon TRF1-TIN2S (arrows, left panels) or TRF1-TIN2L (arrows, right panels) binding to the T270 DNA substrate. XY scale bars: 200 nm. (**E**) Analysis of the TRF1-TIN2S and TRF1-TIN2L mediated DNA loop positions on individual T270 DNA molecules. The loop position was measured from each DNA end (0%) to the loop base with two data points collected from one T270 DNA molecule. T270 only: N=112; TRF1+TIN2S: N=196; TRF1+TIN2L: N=224.

Next, to study TRF1-TIN2 DNA binding at the single-molecule level, we used the linear DNA substrate (5.4 kb) that contains 270 TTAGGG repeats in the middle region (T270 DNA, **Materials and Methods**, **Figure 1A**) (21,49). Previously, AFM and electron microscopy imaging-based studies established that TRF1 specifically binds to the telomeric region and mediates DNA-DNA pairing (21,22,49). To study the function of TIN2, we pre-incubated TRF1, TIN2 (either TIN2S or TIN2L) for 10 minutes, followed by the addition of linear T270 DNA and additional incubation for 10 minutes. The ratio of TRF1:TIN2:DNA was at 1:1:0.17 (300 nM:300 nM:50 nM). AFM imaging of samples deposited onto mica surfaces after dilution (10-fold) revealed heterogeneity in the DNA conformation as well as the size of protein complexes (**Figures 1** and **2**). We categorized the TRF1-TIN2-DNA complexes (N=1283) into three types: individual DNA molecules with a single protein complex bound along the linear DNA contour (21.4% ± 6.2% of total DNA molecules, **Figure 1B**), individual DNA molecules with a loop that is mediated by a protein complex (7.4% ± 4.6% of total DNA molecules, **Figure 1D**), and clusters of multiple T270 DNA fragments bridged by protein complexes (*trans*-interactions, 38.5 ± 3.7% of total DNA molecules, **Figure 2**). In comparison, for TRF1 alone with T270 DNA under the same conditions, complexes with protein-mediated DNA looping and clusters of multiple T270 DNA fragments bridged by proteins were rare (<5%, N=568 DNA molecules). Furthermore, for the non-telomeric DNA (noTel DNA, 4.1 kb) in the presence of TRF1 and TIN2L, the percentage of DNA molecules with bound proteins was lower (19.8%, N=765 DNA molecules, **Figure 2C**) compared to what was observed on the telomeric DNA. On the non-telomeric DNA, clusters of multiple DNA fragments bridged by proteins (6.2% ± 2.1%) were significantly (p<0.05) reduced. These results suggested that TRF1-TIN2-mediated *trans-* and *cis*-interactions are telomeric DNA specific.

**Figure 2.**
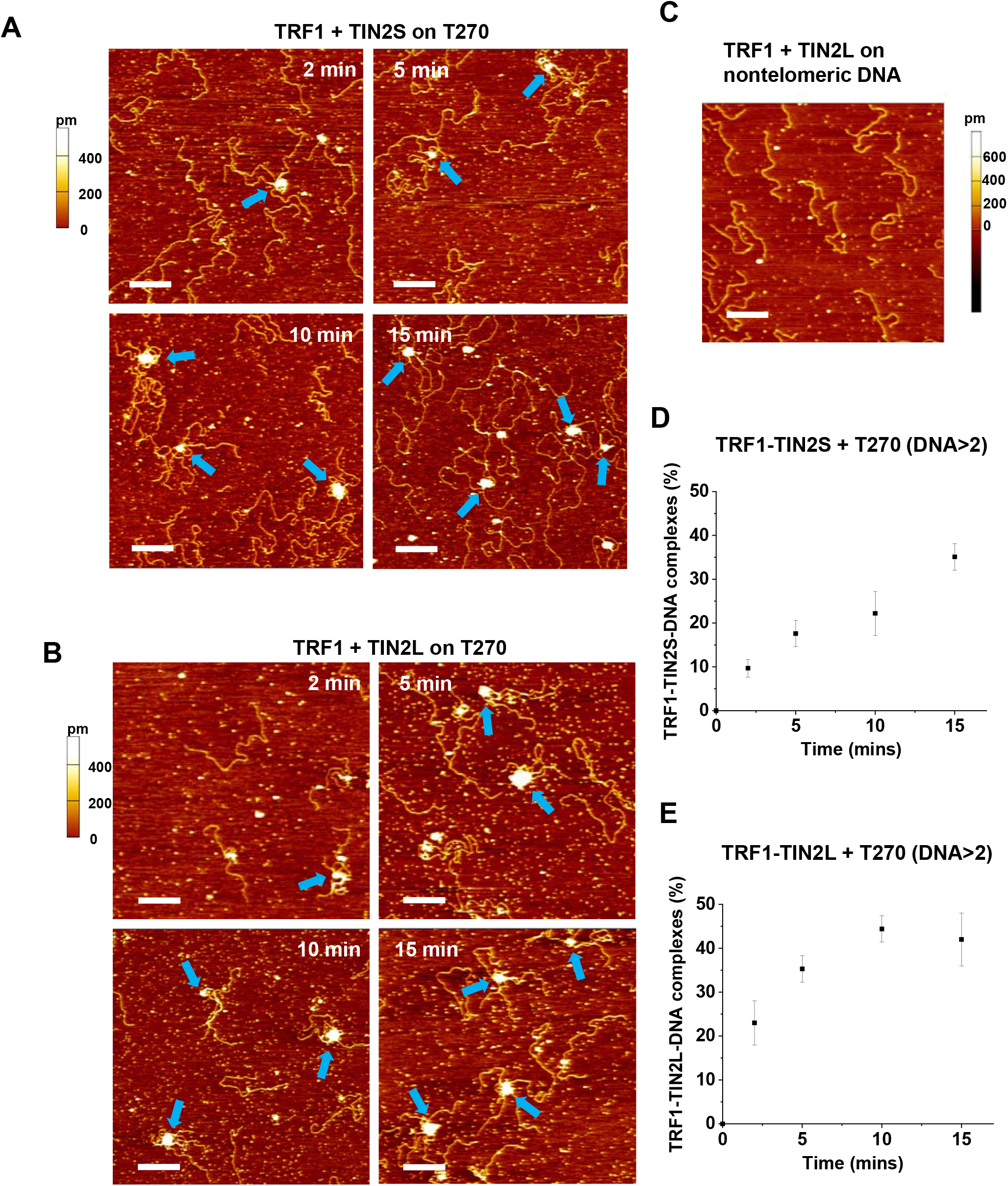
TRF1-TIN2 forms large complexes and bridges multiple dsDNA fragments in a telomeric sequence-dependent manner. (**A** and **B**) Representative AFM images of TRF1-TIN2S (**A**) and TRF1-TIN2L (**B**) complexes showing bridging of multiple fragments of T270 DNA (5.4 kb) after incubation for 2 min, 5 min, 10 min, and 15 min. Arrows point to complexes containing more than two T270 fragments. (**C**) A representative image of TRF1-TIN2L on the control nontelomeric DNA (noTel, 4.1 kb) after incubation for 15 min. XY scale bars: 500 nm. (**D** and **E**) Percentages of complexes containing more than two T270 DNA fragments bridged by TRF1-TIN2S (**D**) or TRF1-TIN2L (**E**) at different incubation times. The percentages of complexes with greater than two DNA fragments were 9.6 ± 2.3% (2 min), 17.6% ± 3.1% (5 min), 22.3% ± 5.2% (10 min), and 34.8% ± 3.5% (15 min) for TRF1-TIN2S, and 23.1% ± 4.6% (2 min), 35.2% ± 2.9% (5 min), 44.1% ± 3.2% (10 min), and 42.5% ± 5.1% (15 min) for TRF1-TIN2L. Each data set was from at least three independent experiments. Error bars: SEM from three independent experiments.

Next, to investigate the DNA binding properties unique to TRF1-TIN2, we identified protein complexes containing TIN2 on DNA. Based on the difference between the AFM height of TRF1 alone (0.48 ± 0.13 nm), we used an AFM height cutoff at 1 nm to select the protein complexes on T270 DNA that contained TIN2. Based on this selection criteria, the majority of the TRF1-TIN2 protein complexes (55% for TIN2S and 75% for TIN2L) were located at the telomeric region (35%-50% from the closest DNA end) on the linear T270 DNA (**Figure S3A**). In addition, the distribution of TRF1-TIN2L complexes on the control linear noTel DNA was random along the DNA contour (**Figure S3B&C**). These results indicated that TRF1-TIN2 specifically binds to the telomeric DNA sequences. Furthermore, to detect whether or not TRF1-TIN2 compacted DNA, we measured the contour length of T270 DNA and the control non-telomeric DNA in the absence or presence of proteins. For TRF1 only, the T270 contour length was shortened slightly from 1.71 μm (± 0.05 μm) for DNA alone to 1.64 μm (± 0.06 μm) (**Figure 1C**). The linear T270 DNA contour length in the presence of TRF1-TIN2 displayed wider distributions and was significantly (p<0.001) further shortened from the level observed with TRF1 only (1.58 μm ± 0.11 μm for TRF1-TIN2S and 1.59 μm ± 0.1 μm for TRF1-TIN2L, **Figure 1C**). In comparison, the lengths of the linear noTel DNA were comparable in the absence or presence of TRF1-TIN2L (**Figure S3D**). In summary, these results established that TRF1 and TIN2 together compact telomeric DNA.

To verify that TRF1-TIN2 complexes mediate DNA looping (*cis*-interactions) specifically at the telomeric region, we quantified the position of the base of protein-mediated DNA loops by directly measuring the length of each DNA arm (**Figure 1E**). We treated 0% to 35% as non-telomeric regions, and 35% as telomeric and non-telomeric region boundaries. TRF1-TIN2 mediated DNA looping either within the telomeric region (11.4% for TIN2S and 13.4% for TIN2L), close to the two boundaries of the telomeric region (28.6% for TIN2S and 21.4% for TIN2L), or between telomeric and non-telomeric regions (60.0% for TIN2S and 65.2% for TIN2L). These results demonstrated that TRF1 and TIN2 together mediated DNA looping in a telomeric sequence-dependent manner.

Furthermore, in addition to the individual DNA molecules with bound proteins, we also observed that TRF1-TIN2S and TRF1-TIN2L formed large complexes that bridged multiple strands of linear T270 DNA molecules in a time-dependent manner (**Figure 2**). Strikingly, after 15 minutes of incubation, a significant percentage of linear T270 DNA molecules (34.8% ± 3.5% for TRF1-TIN2S and 42.5% ± 5.1% for TRF1-TIN2L) were in the protein-DNA clusters with more than two T270 fragments (**Figure 2D&E**). The sizes of protein complexes involving multiple copies of T270 when both TRF1 and TIN2 were present (TRF1-TIN2S: 6816 ± 4788 nm^3^, TRF1-TIN2L: 7286 ± 5229 nm^3^, **Figure S4**) were at least 45 times greater than those of combined volumes of a TRF1 dimer (~105 nm^3^) and TIN2 monomer (~41 nm^3^, **Figure S1**). These results indicate that multi-protein TRF1-TIN2 complexes were present when bridging multiple fragments of T270 DNA. Collectively, results from AFM imaging established that both TIN2S and TIN2L facilitate *cis-interactions* leading to DNA compaction and *trans*-interactions leading to DNA-DNA bridging in a telomeric sequence-dependent manner.

### TIN2 increases TRF1-mediated telomeric DNA-DNA bridging lifetimes

To gain mechanistic insight into how TIN2 influences DNA-DNA bridging by TRF1, we applied single-molecule TIRFM fluorescence imaging to monitor DNA-DNA bridging in real-time. First, Cy5-labeled and biotinylated double-stranded telomeric DNA substrates containing 6 TTAGGG repeats were anchored onto PEGylated quartz slides through biotin-streptavidin interactions (**Materials and Methods**, **Figure 3A**). Then, we introduced TRF1 or TRF1-TIN2 (TIN2S or TIN2L, 100 nM each protein) into the flow cell, followed by washing with the imaging buffer (**Materials and Methods**). After 10 minutes of incubation, we added the telomeric Cy3-DNA with 6 TTAGGG repeats but lacking biotin as free DNA from solution (5 nM, **Materials and Methods**) along with proteins (TRF1 with or without TIN2) into the flow cell. To reveal the dynamics of the protein-mediated colocalization of the Cy5- and Cy3-DNA (*trans*-interactions), we illuminated the slide surface with a red laser, followed by a green laser, to excite the Cy5 and Cy3 fluorophores, respectively, and captured 120-second movies. To monitor whether the free Cy3-DNA from solution was bridged by proteins to the Cy5-DNA anchored onto the surface, the spots that displayed signals in the Cy5 channel were located first. Then, a map was used to locate the spots in the Cy3-DNA channel (**Figure S5A**) and to reveal the temporal dynamics of the colocalization of Cy5- and Cy3-DNA (**Figure 3**).

**Figure 3.**
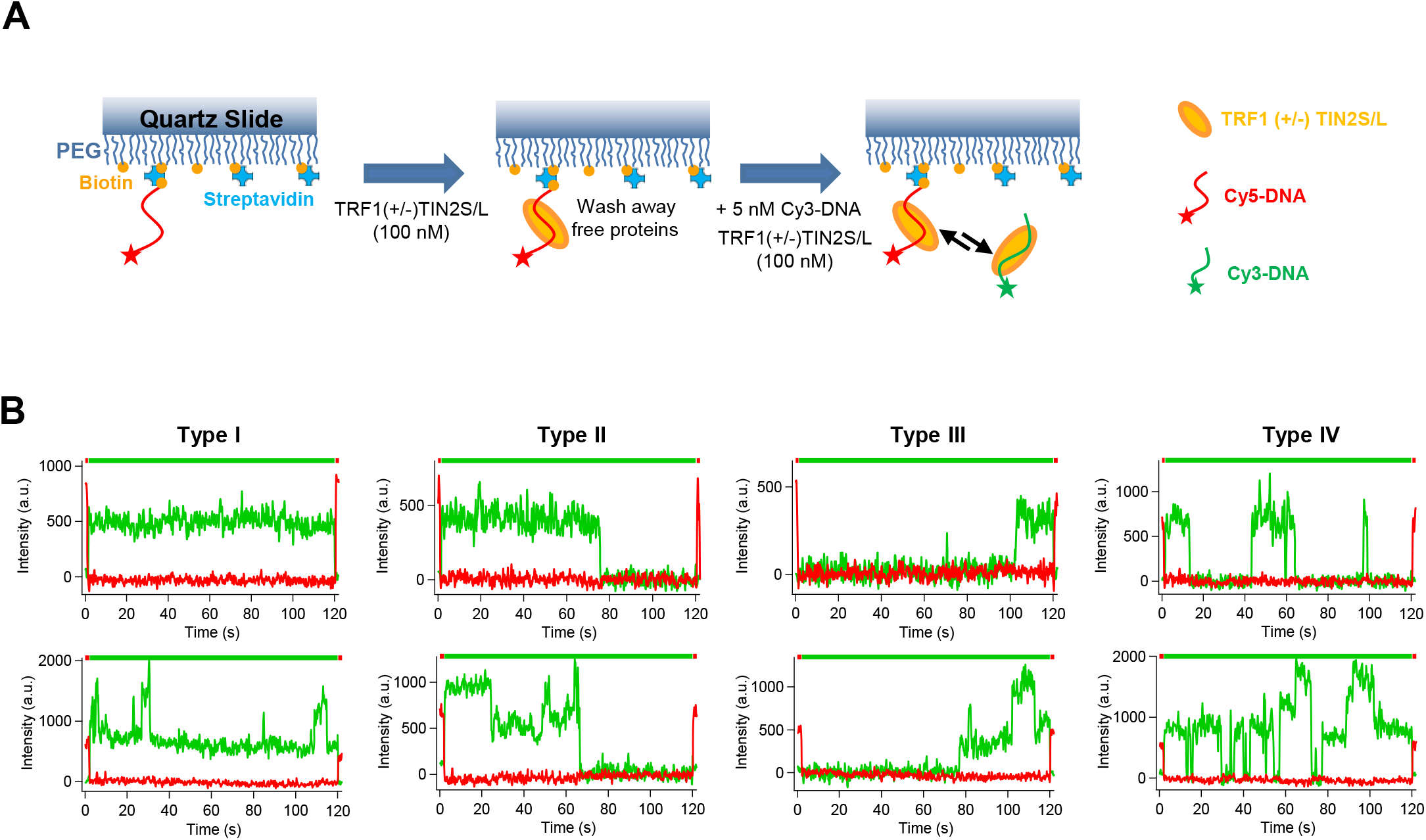
TIN2 facilitates TRF1-mediated bridging of telomeric dsDNA revealed by TIRFM imaging. (**A**) Schematics of the experimental setup for monitoring DNA-DNA bridging (*trans*interactions) mediated by TRF1-TIN2 using TIRFM imaging. (**B**) Four types of single-molecule traces of TRF1-TIN2 mediated DNA-DNA bridging events observed using TIRFM imaging. One or multiple Cy3-DNA molecules were bridged to the surface-anchored Cy5-DNA. The lines on the top indicate laser excitation sequence.

To validate that Cy3- and Cy5-DNA colocalization was mediated by proteins in a telomeric sequence-dependent manner, and not through nonspecific interactions, we conducted a series of control experiments under the same experimental conditions. First, we conducted control experiments without proteins (**Figure S5B**), and found only 1.5% of traces (N=4104) showed colocalization of Cy5- and Cy3-DNA signals. These results demonstrate that there was minimal nonspecific binding of free DNA to the surface without proteins. Furthermore, to rule out the possibility of nonspecific attachment of proteins and Cy3-DNA to the surface, we carried out additional controls without Cy5-DNA on the surface, but with the same concentrations of proteins and Cy3-DNA in the imaging chamber (**Figure S5C**). This control experiment showed that less than 5.5% of traces at randomly selected spots (N=500) displayed Cy3-DNA signals on the surface that simulated experiments using the Cy5-DNA on the surface. We, therefore, concluded there was minimal colocalization of Cy5- and Cy3-DNA signals due to random coincidence on the surface. Next, to test the DNA binding specificity of the TRF1-TIN2 system, we repeated the experiments using the Cy3-DNA with scrambled sequences. After incubating free non-telomeric Cy3-DNA in the imaging chamber, less than 5% of all traces (N=2788 with TRF1-TIN2 and N=3574 without proteins) showed colocalized Cy3- and Cy5-DNA signals (**Figure S5D&E**). In summary, these control experiments demonstrated that under the experimental conditions, the TIRFM imaging platform enables us to investigate TRF1-TIN2 mediated telomeric DNA-DNA bridging (*trans*-interactions) in real-time and at the single-molecule level.

To investigate whether or not TIN2 influences the efficiency and dynamics of TRF1-mediated telomeric DNA-DNA bridging, we directly compared Cy5-Cy3 colocalization signals when only TRF1, or both TRF1 and TIN2 were present. When the telomeric Cy5-DNA was immobilized on the surface, and TRF1 (100 nM) was present in the flow cell, the percentage of Cy5 traces colocalized with telomeric Cy3-DNA signals was 16.6% (N=2369). This percentage increased to over 40% (N=3540) in the presence of both TRF1 and TIN2 (TIN2S or TIN2L). To further interpret the data, we classified the traces with Cy5-Cy3 DNA colocalization signals into four types (**Figure 3B**). Type I: stable bridging in which the Cy3-DNA in solution was bridged to the Cy5-DNA on the surface from the beginning to the end of the movie; Type II: the Cy3-DNA was bridged to the Cy5-DNA at the beginning of the movies and then released (or photobleached); Type III: Cy3-DNA was bridged to Cy5 DNA during the middle of the movies and did not dissociate from each other till the end of the movies; Type IV: multiple transient DNA-DNA bridging and releasing events. It is worth noting that we categorized the traces based on Cy3-DNA signal duration. Based on the Cy3 signal intensities, each category includes events with either one or multiple Cy3-DNA molecules bridged to the surface-anchored Cy5-DNA (**Figure 3B**). When TRF1 (100 nM) alone was present in the chamber, only 1% (± 0.3%) of all event traces showed Type I stable bridging. The dwell time and dissociation time of Cy5-Cy3 DNA bridging events were 1.95 s and 3.38 s, respectively (**Figure S6**). Importantly, the percentage of Type I stable bridging events that lasted throughout the observational window (120 s) increased significantly (p<0.05) to 10.6% (± 5.6%) with the addition of TIN2S (100 nM) and to 19.0% (±10.6%) with the addition of TIN2L (100 nM, **Table 1**). These results demonstrate that with TRF1 only telomeric DNA-DNA bridging events were transient, and TIN2 stabilizes TRF1-mediated DNA-DNA bridging.

**Table 1.** The percentages of each type of telomeric DNA-DNA bridging events with TRF1 alone, or TRF1 with increasing concentrations of TIN2S or TIN2L observed using TIRFM.

| Trace Types | TRF1 (100 nM) |  |  |  |  |  |  |
| --- | --- | --- | --- | --- | --- | --- | --- |
|  | No TIN2 | 25 nM | TIN2S<br>50 nM | 100 nM | 25 nM | TIN2L<br>50 nM | 100 nM |
| Type I | 1.0±0.3 % | 2.6 ±1.3 % | 4.1 ±1.0 % | 10.6±5.6 % | 3.7±1.7 % | 7.0±1.1 % | 19.0±10.6% |
| Type II | 2.0±0.8 % | 3.5 ±1.0 % | 9.0 ±7.0 % | 4.9±0.4 % | 3.1±0.5 % | 6.0±2.3 % | 3.9±2.7 % |
| Type III | 0.5±0.1 % | 0.5±0.25 % | 0.7±0.2 % | 1.6±0.9 % | 0.7±0.4 % | 1.7±1.0 % | 3.4±2.1 % |
| Type IV | 13.3 ±1.8 % | 16.0±9.5% | 21.8±3.8 % | 26.8±6.9 % | 13.4±2.3 % | 30.0±8.6 % | 31.2±11.4% |
| No signal | 83.4±0.5 % | 77.5±9.4 % | 64.4±9.3 % | 56.1±2.6 % | 79.1±2.7 % | 54.9±10.6% | 42.8±1.6 % |

Note: Each data set was based on more than 1000 fluorescence traces collected from three to four independent experiments.

Next, to investigate if the stabilization of TRF1-mediated DNA-DNA bridging is TIN2 concentration-dependent, we quantified the percentages of different types of events at a fixed TRF1 concentration (100 nM), but increasing TIN2 concentrations (25 nM, 50 nM, and 100 nM) (**Table 1**). Results from these experiments showed a clear correlation between increasing TIN2 concentrations and higher percentages of DNA-DNA bridging events. The total percentage of telomeric DNA-DNA bridging signals increased from 22.6% (± 1.6%) at 25 nM to 43.9% (± 2.6%) for TIN2S (100 nM), and from 20.9% (± 0.4%) at 25 nM to 57.5% (± 5.7%) for TIN2L (100 nM). Furthermore, the percentages of Type I stable bridging traces were significantly (p<0.05) increased with higher concentrations of TIN2S (from 2.6% ± 1.3% at 25 nM TIN2S to 10.6% ± 5.6% at 100 nM TIN2S, **Table 1**). A similar trend was observed when the TIN2L concentration was increased (from 3.7% ± 1.7% of Type I at 25 nM TIN2L to 19.0% ± 10.6% at 100 nM TIN2L). These results indicate that TIN2 increases the lifetimes of TRF1 mediated DNA-DNA bridging in a TIN2 concentration-dependent manner.

In addition, consistent with observations from AFM imaging (**Figure 2**), a significant population of traces showed bridging events involving multiple telomeric Cy3-DNA molecules (**Figure 4A&B**). By applying the Chung-Kennedy model (53,54), we classified the traces based on the Cy3 signal intensity levels relative to the first base level in each trace to obtain the number of telomeric Cy3-DNA molecules bridged by proteins (**Figure 4B**). This analysis showed that the percentages of complexes bridging multiple copies of the Cy3-DNA (n>3) increased with higher TIN2 concentrations (**Figure 4C&D**). Specifically, while keeping TRF1 at a fixed concentration (100 nM), increasing TIN2S concentrations (25 nM, 50 nM, and 100 nM) led to a consistent increase in the percentages of telomeric Cy5-Cy3 DNA bridging events involving multiple Cy3-DNA molecules (n>3). Overall, increasing the TIN2S concentration from 25 nM to 100 nM led to an ~4-fold increase (from 7.8% ± 2.1% to 34.7% ± 5.6%) in the percentage of bridging events involving multiple copies of telomeric Cy3-DNA molecules (n>3, **Figure 4C**). Under the same experimental conditions, increasing concentrations of TIN2L led to the same trend. The DNA-DNA bridging events involving multiple copies of telomeric Cy3-DNA molecules (n>3) increased from 10.4% (± 5.3%) at 25 nM TIN2L to 28.9% (± 8.1%) at 100 nM TIN2L (**Figure 4D**). Taken together, results from single-molecule TIRFM imaging experiments demonstrated that TIN2 increases the lifetimes of TRF1-mediated telomeric DNA-DNA bridging events. Furthermore, TRF1-TIN2 complexes are capable of promoting the bridging of multiple copies of telomeric DNA.

**Figure 4.**
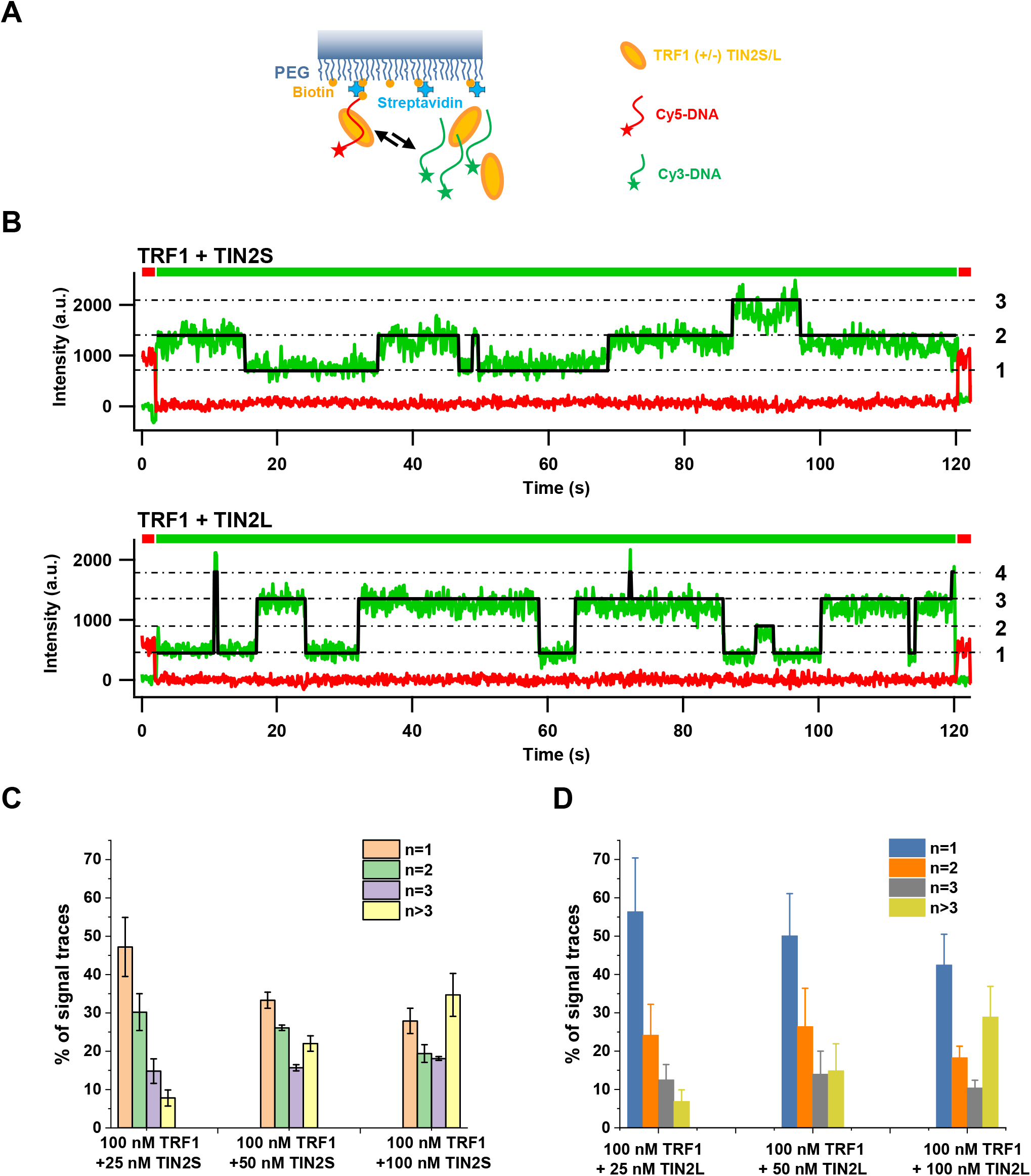
TRF1-TIN2 bridges multiple telomeric Cy3-DNA molecules to surface-anchored telomeric Cy5-DNA. (**A**) Schematics of multiple telomeric Cy3-DNA molecules being bridged to the surface-anchored telomeric Cy5-DNA by the TRF1-TIN2 complex. (**B**) Representative traces from TRIFM imaging showing TRF1-TIN2S (top) and TRF1-TIN2L (bottom) bridged multiple telomeric Cy3-DNA molecules to the surface-anchored telomeric Cy5-DNA. Dashed-dot lines indicate one, two, three, and four telomeric Cy3-DNA molecules being bridged to the telomeric Cy5-DNA based on the fluorescence intensity of the Cy3 signals. The black lines are the fitting from the Chung-Kennedy model. (**C** and **D**) Percentages of telomeric Cy5-Cy3 DNA bridging events with 1, 2, 3, or greater than 3 Cy3-DNA molecules at increasing TIN2S (**C**) or TIN2L (**D**) concentrations. The TRF1 concentration was kept constant (100 nM). Each data set was based on greater than 500 fluorescence traces collected from more than three independent experiments. Error bars: SEM.

### TRF1 loads TIN2 specifically onto the telomeric regions forming stable TRF1-TIN2 complexes

Next, to further investigate the DNA binding specificity and stability of TRF1-TIN2 complexes on longer telomeric DNA substrates, we applied the DNA tightrope assay. This assay (**Materials and Methods**) is based on oblique-angle fluorescence microscopy imaging of QD-labeled proteins on DNA anchored between silica beads (21,55–57). DNA tightropes are formed by stretching DNA between poly-L-lysine coated beads under hydrodynamic flow inside a flow cell. The defined spacing between specific DNA sequences and structures on DNA tightropes enables us to correlate DNA binding events with specific DNA sequences or structures (21,49–51,58). Specifically, to study telomere binding proteins, we ligated linear T270 fragments to form DNA tightropes with telomeric regions (270 TTAGGG repeats) at defined spacing (**Figure 5A**). The lengths of the T270 DNA tightropes are typically in the range of approximately 2.1 to 22 μm, corresponding to tandem ligation of 2-12 of T270 fragments (5.4 kb). To monitor TRF1 and TIN2 (TIN2S or TIN2L) on DNA in real-time, we conjugated His-tagged TRF1 to streptavidin-coated QDs (strep-QDs) through the biotinylated multivalent chelator tris-nitrilotriacetic acid (^BT^tris-NTA) linker, and HA-tagged TIN2 to primary HA antibody-coated QDs (HA-Ab-QDs) (**Materials and Methods**, **Figure 5B**). Previously, we established that QD-labeling of TRF1 using strep-QDs does not significantly affect its telomeric DNA binding specificity (21). Without proteins, there was no significant binding of either strep-QD or Ab-QD alone to DNA tightropes. Furthermore, without TRF1, no significant binding of either TIN2S-QDs or TIN2L-QDs to T270 DNA tightropes was observed. To rule out possibilities of cross-talk between the two QD-labeling strategies, we performed control experiments with combinations of TRF1 with HA-Ab-QDs or TIN2S/TIN2L with strep-QDs. Under these conditions, no significant binding of QDs on DNA tightropes was observed. These control experiments validated the specificity of the two QD labeling strategies.

**Figure 5.**
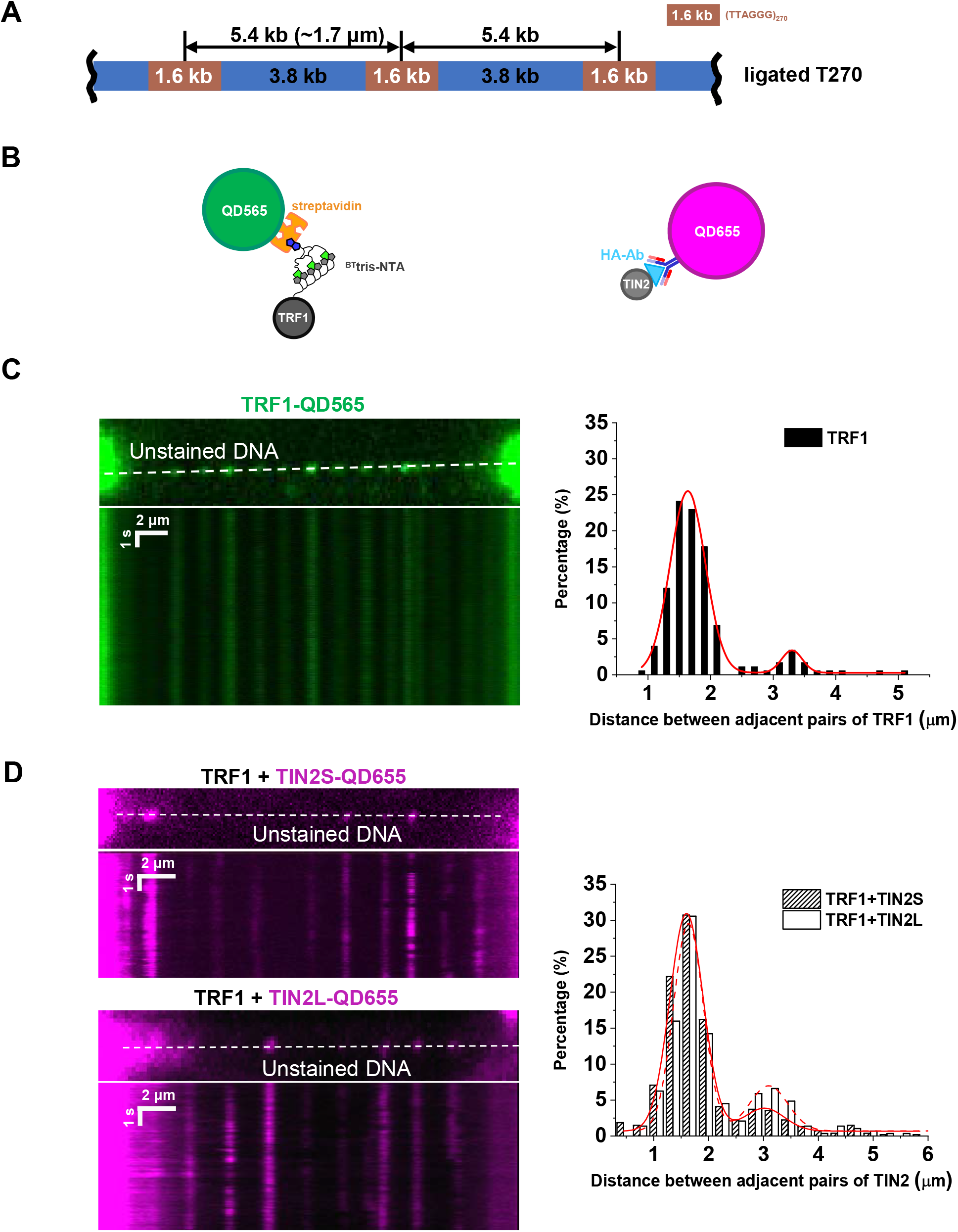
TRF1 loads TIN2 specifically at telomeric regions on T270 DNA tightropes. (**A**) Schematics of the ligated T270 DNA substrate. (**B**) QD labeling schemes for TRF1 and TIN2. His-tagged TRF1 was labeled with strep-QDs through the ^Bt^tris-NTA compound (left panel). HA-tagged TIN2 was conjugated to HA-Ab-QDs (right panel). (**C**) TRF1-QDs on the T270 DNA tightrope. Left panel: representative fluorescence images and kymograph of green (565 nm) strep-QD-labeled TRF1. Right panel: the spacing between two adjacent TRF1 on T270 DNA tightropes. Fitting the data with a double Gaussian function (red line in the right panel) shows peaks centered at 1.63 μm and 3.29 μm (N=185, R^2^> 0.98). (**D**) TIN2-QDs on the DNA tightrope in the presence of unlabeled TRF1. Left panels: representative fluorescence images and kymographs of red (655 nm) HA-Ab-QD-labeled TIN2 (TIN2S: top and TIN2L: bottom). Right panel: the distance between two adjacent TIN2S-QDs (N=538) or TIN2L-QDs (N=288) on T270 DNA tightropes. Fitting the data with double Gaussian functions (R^2^> 0.98) shows peaks centered at 1.60 μm and 3.02 μm for TRF1-TIN2S (solid red line), and 1.63 μm and 3.09 μm for TRF1-TIN2L (dotted-red line).

To establish that TRF1 specifically binds to the telomeric region on DNA tightropes, we measured the pairwise distance between adjacent TRF1-QDs (**Figure 5C**). The spacing was consistent with the distance between telomeric regions (**Figure 5C**). To monitor the recruitment of TIN2 onto DNA tightropes by TRF1, after adding unlabeled TRF1 (10 nM) into the flow cell and 5 minutes of incubation, TIN2-QDs (either TIN2S or TIN2L, 10 nM) were introduced into the flow cell using a syringe pump. Then the buffer flow was shut off to enable freely diffusing TIN2-QDs in solution to bind TRF1 on DNA tightropes. We observed long-lived TIN2S-QDs and TIN2L-QDs on T270 tightropes, with ~97% of TIN2-QDs remaining on DNA tightropes till the end of the observational windows (2 minutes: N=1043; 5 minutes: N=167). Furthermore, the majority of TIN2-QDs loaded by TRF1 (97%, N=167) onto T270 DNA tightropes were static throughout the observational window (5 minutes). For both TIN2S and TIN2L, the histograms of the pairwise distance between adjacent binders displayed two peaks (**Figure 5D**). The two peaks were centered at 1.60 μm and 3.02 μm for TIN2S (N=538), and 1.63 μm and 3.09 μm for TIN2L (N=288). Thus, the spacing of TIN2S and TIN2L were consistent with the expected spacing between telomeric regions on the T270 tightropes (**Figure 5A**).

Furthermore, to directly visualize the interaction between TRF1 and TIN2 on DNA tightropes, His-TRF1 and HA-TIN2 were differentially dual-color labeled (**Figure 5B, Figure S7**). Green (565 nm) TRF1-QDs (10 nM) and red (655 nm) TIN2-QDs (10 nM) were premixed for 10 minutes before being introduced into the flow cell. Under these experimental conditions, 36.0% of the total protein-QDs on T270 DNA tightropes were dual-color labeled (N=339). Importantly, the spacing of dual-color labeled TRF1-TIN2S complexes was also consistent with the expected distance between telomeric regions on T270 tightropes (**Figure S7**). In summary, these results showed that TIN2S and TIN2L are recruited by TRF1 specifically to the telomeric regions. Furthermore, TRF1-TIN2 form stable and long-lasting complexes on telomeric DNA.

### TIN2S and TIN2L promote stable bridging of long telomeric DNA from solution to DNA tightropes

Having established that TRF1 loads TIN2 specifically at telomeric regions on DNA tightropes, we went further to test whether TIN2 facilitates TRF1-mediated bridging of physiologically relevant long telomeric DNA molecules. For these experiments, the telomeric DNA strands anchored in place were T270 DNA tightropes between micron-sized silica beads. The free DNA strands introduced into the flow cell were biotinylated T270 fragments (270 TTAGG repeats without the flanking non-telomeric regions, ^BT^pT270) or the control non-telomeric DNA (^BT^noTel, **Materials and Methods**, **Figure 6A**). To visualize the ^BT^pT270 DNA and ^BT^noTel, we labeled the DNA strands with red (655 nm) strep-QDs. AFM imaging validated that 60.7% (± 3.1%) of the ^BT^pT270 and 67.7% (± 6.7%) of the ^BT^noTel DNA strands were labeled with strep-QDs (**Figure S8**). With ^BT^pT270-QDs alone, or ^BT^pT270-QDs along with TIN2S or TIN2L in the flow cell, we did not observe any ^BT^pT270-QD signals on T270 DNA tightropes. Thus, we concluded that, in this experimental setup, since ^BT^pT270 does not nonspecifically pair with DNA tightropes, localization of QD signals on DNA tightropes would provide a direct read-out of bridging of ^BT^pT270-QDs to DNA tightropes by proteins.

**Figure 6.**
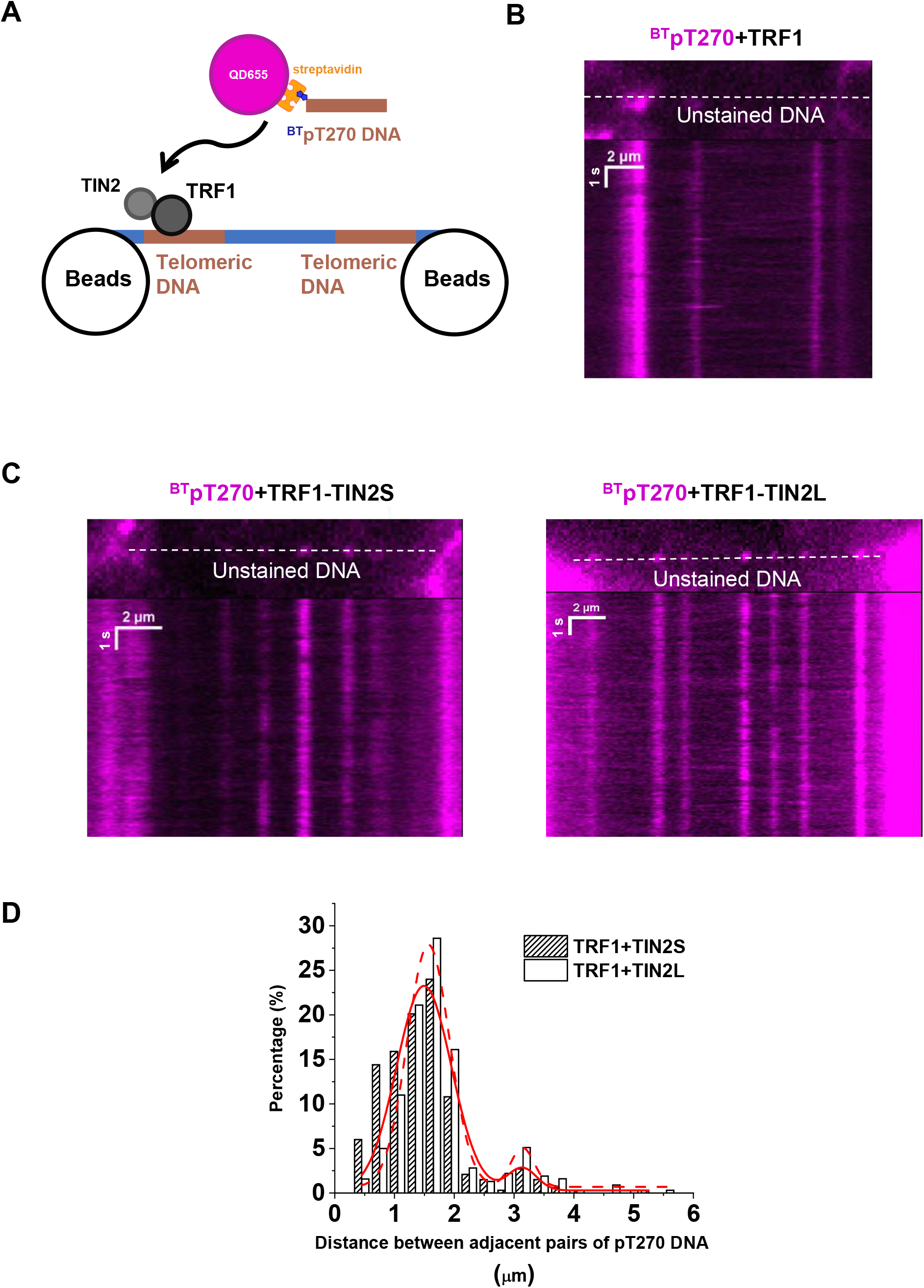
TIN2 facilities TRF1-mediated bridging between the telomeric DNA from solution and T270 tightropes. (**A**) Schematics of the TRF1-TIN2 complex bridging QD-labeled ^BT^pT270 DNA to the T270 DNA tightrope. Brown: telomeric sequences. Blue: nontelomeric sequences. (**B**) Representative fluorescence image and kymograph showing red (655 nm) QD-labeled ^BT^pT270 being bridged to the T270 tightrope in the presence of unlabeled TRF1. (**C**) Representative fluorescence images and kymographs showing red (655 nm) QD labeled ^BT^pT270 being bridged to T270 tightropes in the presence of unlabeled TRF1 and TIN2 (left panel: TIN2S; right panel: TIN2L). (**D**) The distance between two adjacent ^BT^pT270 signals on T270 DNA tightropes. Fitting the data with double Gaussian functions (R^2^>0.86) shows peaks centered at 1.49 μm and 3.13 μm for TRF1-TIN2S (solid red line, N=321 ^BT^pT270-QD signals), and 1.57 μm and 3.14 μm for TRF1-TIN2L (dotted-red line, N=318 ^BT^pT270-QD signals, R^2^>0.95).

To assess whether or not TIN2 facilitates TRF1-mediated DNA-DNA bridging, we added unlabeled TRF1 (10 nM) without or with TIN2 (TIN2S or TIN2L, 10 nM) into the flow cell and incubated for 5 minutes to allow proteins binding to T270 DNA tightropes. Then we introduced ^BT^pT270-QDs (5 nM) along with additional TRF1 (10 nM), either without or with TIN2 (10 nM). While ^BT^pT270-QDs on DNA tightropes were sparse when only TRF1 was present (1.86 ± 0.98 QD signals per 10 μm of DNA, **Figure 6B**), higher densities of ^BT^pT270 fragments were bridged to T270 DNA tightropes when both TRF1 and TIN2 were present (5.71 ± 1.55 QD signals per 10 μm of DNA for TIN2S and 5.15 ± 1.43 QD signals per 10 μm of DNA for TIN2L, **Figure 6C**). Furthermore, with control non-telomeric DNA (^BT^noTel), the numbers of DNA molecules bridged to T270 tightropes were significantly (p<0.05) lower (1.39 ± 0.78 per 10 μm for TRF 1,2.02 ± 0.89 per 10 μm for TRF1 with TIN2S, and 1.98 ± 0.08 per 10 μm for TRF1 with TIN2L). It is worth noting that since we did not stain DNA tightropes and empty DNA tightropes without bridged DNA strands were not counted, this measurement could underestimate the difference in the bridging density between different conditions. ^BT^pT270-DNA tightrope bridging events were long-lasting with 95.5% (± 3.1%, N=438) of the ^BT^pT270 DNA molecules staying on DNA tightropes till the end of the 5-minute observational windows. The spacing between TRF1-TIN2 mediated DNA bridging events on T270 tightropes was centered at 1.49 μm and 3.13 μm for TIN2S (N=321), and 1.57 μm and 3.14 μm for TIN2L (N=318, **Figure 6D**). Thus, the spacing between ^BT^pT270 signals on the DNA tightropes was consistent with the distance between telomeric regions (**Figure 5A**). Considering an average center-to-center spacing of 1.5 μm between (TTAGGG)270 regions on DNA tightropes, 6 (TTAGGG)270 regions are expected for every 10 μm of DNA tightropes. An occupation of ~5 ^BT^pT270 signals per 10 μm of DNA tightropes suggested that, in the presence of TRF1-TIN2, the majority of (TTAGGG)270 regions contained protein bridged ^BT^pT270 DNA.

To further confirm that TIN2 was present in the DNA-DNA bridging complex, we dual-color differentially labeled HA-tagged TIN2 and ^BT^pT270 with HA-Ab-QDs (red) and strep-QDs (green), respectively (**Figure S9A**). The majority of ^BT^pT270 (~87%, N=28) bridged to T270 DNA tightropes was colocalized with TIN2 (**Figure S9B&C**). Taken together, these results from the DNA tightrope assay established that TIN2 facilitates stable TRF1-mediated bridging of physiologically relevant long telomeric DNA molecules.

### TIN2 protects TRF1-mediated DNA-DNA bridging in the presence of Tankyrase 1

Telomeric DNA binding by TRF1 is regulated by Tankyrase 1. Tankyrase 1, a member of the poly(ADP-ribose) polymerase (PARPs) family, directly interacts with the acidic domain of TRF1 (37,59). Poly(ADP-Ribosyl)lation (PARylation) of TRF1 inhibits telomeric DNA binding by TRF1 i*n vitro* (37). Consistent with these previous observations, Tankyrase 1 at low concentrations predominantly reduced the formation of higher-order TRF1-DNA complexes (Complex III, **Figure S10**), while at a higher concentration, it inhibited the formation of both TRF1-DNA complexes II and III. Previously, it was shown that TIN2-TRF1-Tankyrase 1 form a ternary complex, and TIN2 blocks the modification of TRF1 by Tankyrase 1 *in vitro* (60). Pre-incubation of TRF1 and TIN2S protected TRF1 telomeric DNA binding (compare lanes 6 and 7 in **Figure S10**). To further investigate whether TIN2 protects TRF1-mediated DNA-DNA bridging from Tankyrase 1, we monitored the bridging of QD-labeled ^BT^pT270 DNA on T270 DNA tightropes in the same flow chamber under different conditions. Recording the positions of DNA tightropes enabled us to go back to the same sets of DNA tightropes, and washing the flow chamber extensively with the buffer allowed us to remove proteins and ^BT^pT270 DNA on tightropes before introducing fresh proteins. This imaging scheme enabled sequential experiments in the same flow chamber and direct comparison of protein-mediated bridging of ^BT^pT270 DNA onto T270 tightropes without or with Tankyrase 1 and NAD+ (Tankyrase 1:150 μg/ml; NAD+:120 μM, **Figure 7**).

**Figure 7.**
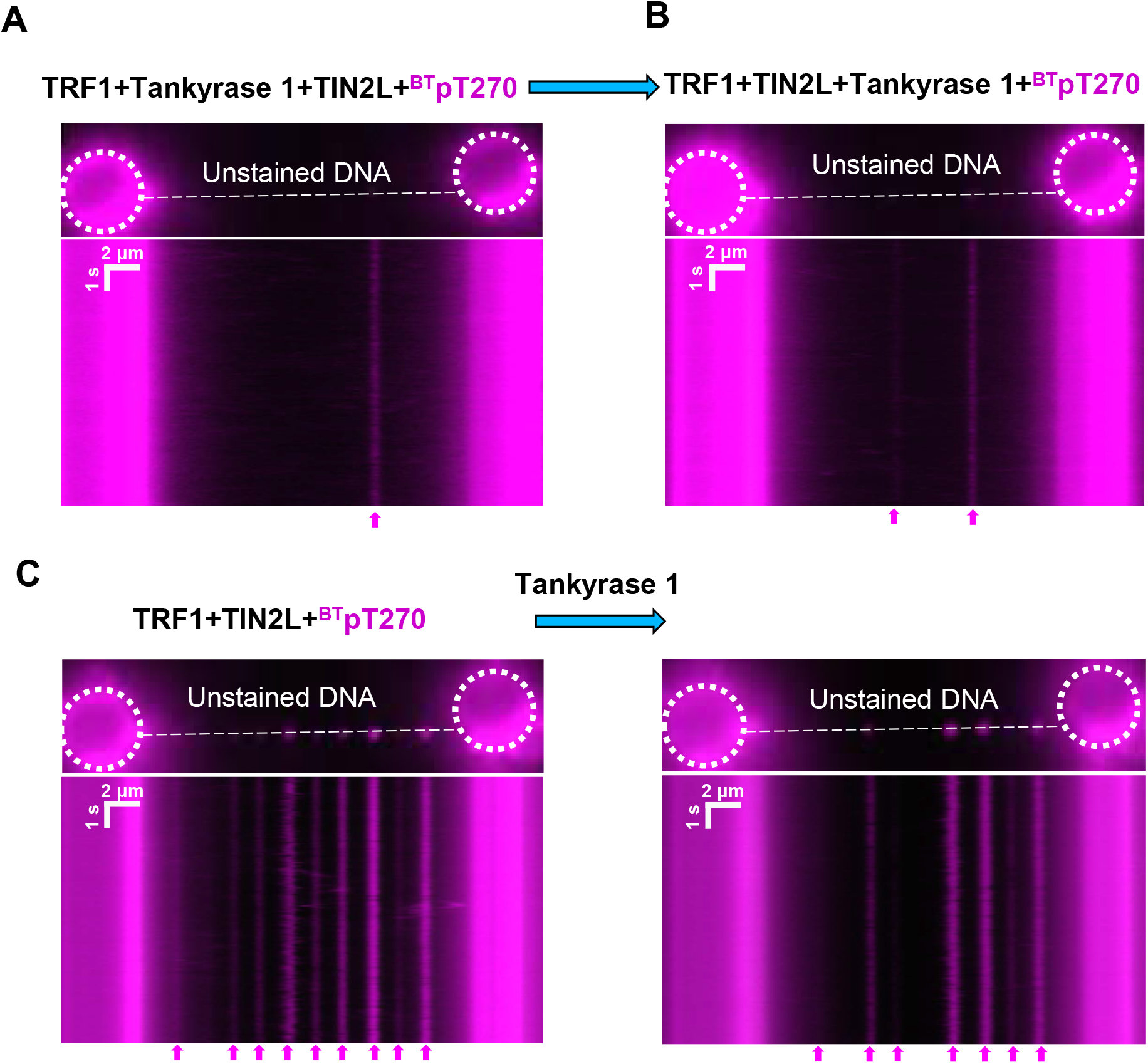
PARylation of TRF1 by Tankyrase 1 reduces DNA-DNA bridging and TIN2 protects TRF1-mediated DNA-DNA bridging in the presence of Tankyrase 1. Examples of fluorescence images (top panels) and kymographs (bottom panels) from four sequential experiments carried out in the same flow chamber to monitor the amount of QD-labeled ^BT^pT270 DNA bridged onto T270 tightropes by TRF1-TIN2L (200 nM each) without or with Tankyrase 1 (150 μg/mL) and NAD+ (120 μM). The following protein solutions were mixed with the QD-labeled ^**BT**^pT270 DNA (200 nM) and diluted 50X using the imaging buffer before being introduced into the flow cell. (**A**) TRF1 was pre-incubated with Tankyrase 1 for 30 mins before the addition of TIN2L and further incubation for 20 mins. (**B**) TRF1 and TIN2L were preincubated together for 20 mins before the addition of Tankyrase 1 and further incubation for 30 mins. (**C**) Left panel: TRF1 and TIN2L were incubated together for 20 mins without Tankyrase 1 and NAD+. Right panel: After recording the TRF1+TIN2L+ ^**BT**^pT270 experiments (left panel) and flushing the flow chamber with the imaging buffer, Tankyrase 1 and NAD+ were introduced into the flow chamber and incubated for 30 mins before recording. The arrows at the bottom of each kymograph point to the QD-labeled ^BT^pT270 DNA bridged onto T270 tightropes.

The first protein mixture contained TRF1 (200 nM) pre-incubated with Tankyrase 1 and NAD+ followed by the addition of TIN2L (200 nM, **Figure 7A**). Next, after washing the flow chamber, we introduced a second protein mixture containing TRF1 (200 nM) pre-incubated with TIN2L (200 nM) before the addition of Tankyrase I and NAD+ (**Figure 7B**). After these two experimental conditions containing Tankyrase 1 and washing the flow chamber, we introduced TRF1 (200 nM) and TIN2L (200 nM) into the flow chamber along with QD-labeled ^BT^T270 DNA fragments to benchmark the level of ^BT^pT270 DNA bridged onto T270 DNA tightropes without Tankyrase 1 and NAD+ (left panel of **Figure 7C**). Finally, we introduced Tankyrase 1 and NAD+ into the flow chamber to monitor the dismantling of TRF1-TIN2L mediated bridging of ^BT^pT270 DNA by Tankyrase 1. To monitor bridging of ^BT^pT270 to DNA tightropes, each protein mixture was incubated with QD-labeled ^BT^pT270 (200 nM) and diluted 50X in the imaging buffer before being sequentially introduced into the flow chamber containing T270 tightropes.

Setting the level of QD-labeled ^BT^T270 observed on T270 tightropes in the presence of TRF1-TIN2L without Tankyrase 1 at 100% (**Table 2**), pre-incubation of TRF1 and Tankyrase 1 before the addition of TIN2L significantly reduced the number of ^BT^T270 DNA bridging events (8.3% ± 1.3%, **Figure 7A** and **Table 2**). In comparison, the number of ^BT^T270 DNA bridging events when TRF1-TIN2L was incubated first, followed by the addition of Tankyrase 1, was higher (16.9% ± 1.1%, **Figure 7B** and **Table 2**). These results support the notion that TIN2 protects TRF1 from modification by Tankyrase 1, and consequently promotes telomeric DNA-DNA bridging by TRF1. Finally, after TRF1-TIN2L mediated DNA-DNA bridging was formed (left panel, **Figure 7C**), the introduction of Tankyrase 1 and NAD+ reduced the ^BT^T270 DNA bridging events (85.8% ± 4.1%, **Figure 7C right panel** and **Table 2**). Importantly, this level of DNA-DNA bridging events was higher than when TRF1-TIN2L-Tankyrase 1 were pre-incubated together before the inclusion of QD-labeled ^BT^T270 DNA fragments (16.9% ± 1.1%, **Table 2**). Furthermore, inhibition of TRF1-TIN2 mediated DNA-DNA bridging on DNA tightropes depends on the presence of both Tankyrase 1 and NAD+ (**Figure S11**). In summary, these results strongly demonstrated that TIN2 is more efficient in protecting TRF1 from the Tankyrase 1 modification after forming large multi-protein TRF1-TIN2-DNA-DNA bridging complexes than its action on free TRF1 without DNA.

**Table 2.**
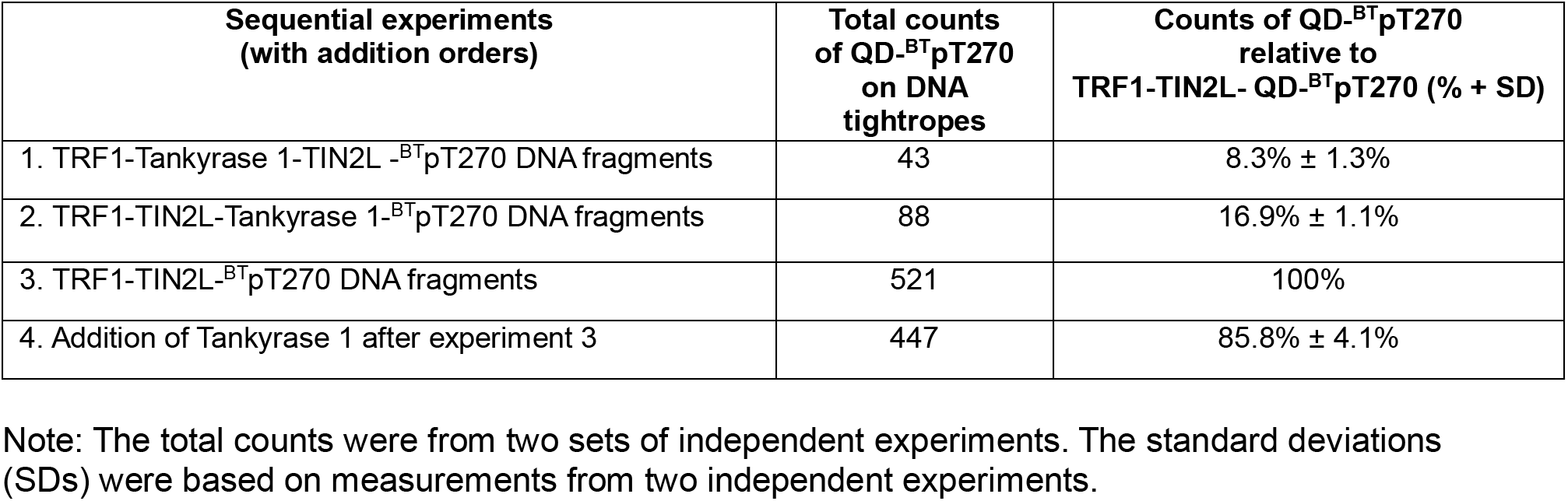
Summary of the level of QD-^BT^pT270 fragments bridged onto T270 tightropes during four sequential experiments in the same imaging chambers.

## DISCUSSION

Despite the central role that TIN2 plays in telomere maintenance, the mechanism underlying its function had been largely unknown. Our results from three single-molecule imaging platforms provide new mechanistic insight supporting the notion that TIN2 facilitates TRF1-mediate DNA compaction and DNA-DNA bridging through several key observations. We discovered that, in the presence of both TRF1 and TIN2, large clusters of protein complexes (AFM volume>500 nm^3^) formed at telomeric DNA regions and shortened the lengths of individual T270 DNA molecules. These results support the notion that multi-protein TRF1-TIN2 complexes facilitate *cis*-interactions on the same telomeric DNA. Furthermore, TRF1-TIN2 promotes *trans*-interactions by stabilizing the bridging of telomeric DNA. AFM imaging revealed the time-dependent formation of clusters of telomeric DNA fragments bridged by multi-protein TRF1-TIN2 complexes. TIRFM tracking of colocalization of surface-anchored telomeric Cy5-DNA and Cy3-DNA from solution showed that *trans*-interactions (DNA-DNA bridging) were telomeric sequence- and TIN2 concentration-dependent. Increasing TIN2 concentrations led to larger percentages of stable telomeric DNA-DNA bridging events (> 2 minutes) and multiple telomeric DNA fragments (n>3) bridged by proteins. Importantly, TRF1-TIN2 promotes stable *trans*-interactions on physiologically relevant long telomeric DNA. With 6 TTAGGG repeats, the percentage of surface anchored telomeric with colocalized Cy3-DNA was 43.9% and 57.2%, for TRF1-TIN2S and TRF1-TIN2L, respectively. With longer telomeric DNA length (270 TTAGGG repeats), the majority (~ 5 out of 6) of the telomeric regions contained telomeric ^BT^pT270 fragments bridged to DNA tightropes. The DNA-DNA bridging events (*trans*-interactions) on T270 DNA tightropes mediated by TRF1-TIN2 were stable and lasted for more than 5 minutes.

The current literature remains inconclusive regarding the activities of shelterin proteins to compact double-stranded telomeric DNA. Previous EM and AFM studies revealed that TRF1 promotes telomeric DNA-DNA pairing and TRF2 compacts telomeric DNA (14,17,20,22,47,49). These results suggested that shelterin proteins might play keys roles in modulating doublestranded telomeric DNA configurations. Furthermore, Bandaria *et al.* measured the telomere size using super-resolution photoactivated localization microscopy (PALM) imaging of mEos2-labeled shelterin proteins and fluorescence *in situ* hybridization-stochastic optical construction microscopy (FISH-STORM) imaging (61). This study suggested that telomeres form compact globular structures through a dense network of specific protein-protein and protein-DNA interactions. Based on this study, TRF1, TRF2, and TIN2 are the main contributors to the telomere compaction, with depletion of TRF1 and TIN2 using siRNA leading to an 8-fold and 6-fold increase in the telomere volume, respectively (61). These results support the notion that TRF1-TIN2 interactions promote telomere compaction. Based on these observations, shelterin-mediated telomere compaction was proposed to be a general mechanism for suppressing DDR pathways at telomeres. However, two recent studies using STORM imaging of telomeres marked with telomeric FISH probes suggested that telomeric compaction upon shelterin removal is either minor or only affects small subsets of telomeres (62,63). When both TRF1 and TRF2 were deleted, which removed the whole shelterin complex from double-stranded regions of telomeres, an average of ~50% increase in the radius of gyration (Rg) and ~157% increase in the convex hull volume of telomeres were observed. Furthermore, other factors such as 53BP1-mediated clustering of dysfunctional telomeres when shelterin was removed from telomeres might contribute to the increase in the Rg/convex hull volume of telomeres. In addition, the telomeres showing DDR did not significantly differ in sizes compared to the ones that were DDR-negative (63). Consistent with these two later studies, single-molecule fluorescence imaging of shelterin proteins on DNA curtains suggested that shelterin binds to telomere DNA as individual complexes (64). In this study, shelterin protein-mediated DNA-DNA bridging was assessed by the retention of the single-tethered DNA onto the double-tethered DNA curtains after buffer flow was turned off. Through using this imaging platform, the authors observed that when shelterin proteins were present, single- and double-tethered DNA molecules separated immediately after buffer flow was switched off. Based on these observations, the authors concluded that shelterin proteins do not promote strong *trans-*interactions.

The discrepancy between results from this previous single-molecule study and our current one could stem from two sources: 1) lengths of the telomeric DNA sequences (32 repeats in the previous study versus 270 repeats in our current study); and 2) freedom of 3D diffusion for at least one telomeric DNA strand that enables the search process, which is critical for protein-mediated DNA-DNA bridging. In the DNA curtain experiment, under flow-stretched configuration, the single-tethered molecules might not be able to fully search in 3D space to pair with the double-tethered DNA. In comparison, in our DNA tightrope assay and TIRFM colocalization setup, the ^BT^pT270-TRF1-TIN2 and Cy3-DNA-TRF1-TIN2 complexes introduced into the flow cell were free to diffuse in 3D. These experimental conditions enabled 3D diffusion to facilitate the bridging of two DNA molecules.

Our results are consistent with a model in which multiple factors contribute to the compaction of telomeres. Compaction of telomeres by additional factors such as nucleosomes or other proteins that remain unidentified could minimize the decompaction effect upon shelterin removal. Furthermore, the results from our study are consistent with the critical role that TIN2 plays in TRF1-TIN2-SA1 mediated sister telomere cohesion that is largely cohesin-ring independent (8,36,65). While siRNA depletion of TRF1 does not significantly affect sister telomere cohesion, depletion of TIN2, which is known to induce degradation of TRF1, leads to a sister cohesion defect. Previously, we reported that SA1 and TRF1 together promote telomeric DNA-DNA pairing tracks (49). The results from AFM, TIRFM imaging, and the DNA tightrope assay reported in this study demonstrated that TIN2 further facilitates TRF1-mediated *trans*-interactions between different telomeric DNA fragments. Furthermore, TIN2 is more efficient in protecting TRF1 from Tankyrase 1 modification when present in the large multi-protein TRF1-TIN2-DNA complexes than when TRF1 is free in solution. Based on results from our current study and previous ones (8,36,49,65), we propose that TIN2 is a key architectural protein that links a telomeric DNA binding protein (TRF1) and a cohesin subunit (SA1) to promote *trans*-interactions that are essential for sister telomere cohesion.

It is worth noting that our single-molecule experiments demonstrate that both TIN2S and TIN2L isoforms show similar activities in promoting TRF1-mediated DNA compaction and DNA-DNA bridging. In order to advance our understanding of the unique biological functions of each TIN2 isoform, future work is needed to uncover differences in their interactions with other protein partners.

## MATERIALS AND METHODS

### Protein purification

Recombinant N-terminal His6-tagged TRF1 was purified using a baculovirus/insect cell expression system and an AKTA Explorer FPLC (GE Healthcare) as described in a previous study (66). N-terminal HA-tagged TIN2L (HA-TIN2L, 1-451 aa) and TIN2S (HA-TIN2S, 1-354 aa) were expressed in the Sf-900 insect cells using the pFastBac1 expression system (GenScript). HA-TIN2L and HA-TIN2S were purified using anti-HA resin and stored in a buffer containing 50 mM Tris-HCl (pH 7.5), 150 mM NaCl, 10% glycerol, 1% NP40, and 1 mM EDTA. The concentrations of TIN2 proteins were determined by the BCA^TM^ protein assay with BSA as the standard (ThermoFisher). The identities of purified HA-TIN2S and HA-TIN2L proteins were confirmed by the Western Blot analysis using the HA antibody (GenScript A00168), and MALDI-TOF mass spectrometry analysis (UNC-Chapel Hill Proteomics Center). His-tagged Tankyrase 1 was purified from SF21 insect cells, as described previously (37).

### DNA substrates

The plasmid containing 1.6 kb telomeric TTAGGG repeats with a 23-bp interruption linking two (TTAGGG) 135 regions (T270, 5.4 kb) was purchased from Addgene (plasmid pSXneo(T2AG3), #12403) (67). For AFM imaging, T270 plasmid was digested by HpaI at 37°C for 4 hr in the CutSmart buffer (New England BioLabs) to place the (TTAGGG)270 at the middle region of the linearized substrate. In order to generate longer DNA substrates for fluorescence imaging, the linearized T270 was ligated using the Quick Ligation^TM^ Kit (New England BioLabs) at room temperature for 1 hr followed by additional incubation at 4°C for overnight. Ligated T270 was purified to remove ligase by phenol-chloroform extraction using the Phase Lock Gel^TM^ (Quantabio). The biotinylated T270 substrate (^BT^pT270) and control DNA (^BT^noTel) were generated through biotinylation of the gel-purified T270 fragment containing the (TTAGGG)270 region (pT270, 1.6 kb) or control nontelomeric DNA (noTel, 4.1 kb) using the 5’ EndTag Labeling DNA/RNA Kit (Vector Laboratories). The biotinylation of the pT270 and noTel fragments was verified using AFM imaging of DNA samples in the presence of streptavidin-coated quantum dots (strep-QDs, Invitrogen). All oligos were purchased from IDT. Cy5/biotinylated DNA (69 bp) with 6 TTAGGG repeats used for the single-molecule TIRF experiments was generated by annealing the Cy5-labeled oligo (Cy5-5’TCTGTAGTGTA *TTAGGGTTAGGGTTAGGGTTAGGGTTAGGGTTAGGGATGTAGGTATGTCACAGCATGA3*’) and complementary biotin-labeled oligo (biotin-5’TCATGCTGTGACATACCTACAT *CCCTAACCCTAACCCTAACCCTAACCCTAACCCTAA*TACACTACAGA3’). Cy3-DNA without biotin (59 bp) was prepared by annealing Cy3-labeled oligo (Cy3-5’ACATACCTACAT*CCCTAA CCCTAACCCTAACCCTAACCCTAACCCTAA*TACACTACAGA3’) and complementary oligo (5’ *TCTGTAGTGTATTAGGGTTAGGGTTAGGGTTAGGGTTAGGGTTAGGGATGTAGGTATGT3*’). The telomeric DNA substrate used for electrophoresis mobility shift assays (EMSAs) was constructed by annealing a 5’ Alexa488-labeled oligo (5’*TTAGGGTTAGGGTTAGGG*ATGTCCAGCAAGCCAGAATTCGGCAGCGTA3’) with its complementary oligo.

### Electrophoresis mobility shift assays (EMSAs)

TRF1 and TIN2 DNA binding reactions were carried out in a buffer containing 20 mM HEPES (pH7.9), 100 mM NaCl, 1mM EDTA, 0.5 mM DTT, 0.25% NP-40, and 5% glycerol. The samples were loaded onto a 5% 29:1 (bisacrylamide:acrylamide) native gel. Electrophoresis was carried out at 150 V for 40 min in 1X TBE buffer at 4°C. The gel was visualized using a Typhoon Phosphorimager (FLA 7000).

### Protein-quantum dot conjugation

Green (565 nm) streptavidin-coated quantum dots (strep-QDs, Life Technologies) were conjugated to N-terminal His6-tagged TRF1 proteins through the multivalent tris-nitrilotriacetic acid chelator (^BT^tris-NTA) as previously reported (21). The primary HA antibody (Abcam) was conjugated to red (655 nm) QDs using the SiteClick^TM^ QD antibody labeling kit (Life Technologies) according to the manufacturer’s standard protocols. The final concentration of HA-Ab-QDs was measured on Nanodrop. HA-Ab-QDs were then conjugated to HA-tagged TIN2S or TIN2L by incubation at a 1:1 ratio for 20 min at room temperature.

### Atomic force microscopy (AFM) imaging

TRF1 and TIN2 were incubated in a buffer containing 20 mM HEPES (pH7.9), 100 mM NaCl and 1mM EDTA at a 1:1 ratio (300 nM:300 nM). After 10 min of incubation at room temperature, linearized telomeric T270 DNA (5.4 kb, 5 nM) or control DNA (noTel DNA, 4.1 kb) was added into the protein mixture and incubated for another 10 min. The sample was diluted 10-fold using the AFM imaging buffer (25 mM HEPES, pH7.5, 25 mM NaOAc, and 10 mM Mg(OAc)2) before being deposited onto a freshly peeled mica surface (SPI supplies).

### The DNA tightrope assay

The oblique angle TIRFM-based DNA tightrope assay was described previously (49,50,55–57). Briefly, we immobilized poly-L-lysine (2.5 mg/mL, M.W.>30000 KDa, Wako Chemicals) treated silica beads onto a PEGlyated coverslip surface, and then introduced ligated DNA substrates into the flow cell using a syringe pump at a flow rate of 300 μL/min to stretch the DNA between treated beads. All protein-QD samples were diluted 100-fold using the imaging buffer (20 mM HEPES, pH7.9, 100 mM NaCl, 0.3 mM MgCl_2_, 1mM EDTA, 0.5 mM DTT and 1 mg/mL BSA) to a final concentration of 10 nM proteins in the imaging chamber. To prevent nonspecific binding, 1X Blocking Reagent (Sigma, catalog number 11096176001) was used in the imaging buffer. For experiments investigating the impact of Tankyrase 1 on the efficiency TRF1-TIN2 mediated bridging of QD-labeled linear telomeric ^BT^pT270 DNA to T270 DNA tightropes, we carried out a series of four sequential experiments in the same flow chambers. After videos were taken under each condition, the flow chamber was flushed with the imaging buffer (200 μL, ~6X of the flow chamber volumes) to set up for the next experiment. We confirmed that greater than 90% of the QD-^BT^pT270 on DNA tightropes were washed off under this condition. The four sequential experiments were designed as follows: 1) TRF1 (200 nM) was pre-incubated with Tankyrase 1 (150 μg/mL) and NAD+ (120 μM, MilliporeSigma, catalog number 481911) for 30 mins before the addition of TIN2L (200 nM) and further incubation for 20 mins. 2) TRF1 (200 nM) and TIN2L (200 nM) were pre-incubated together for 20 mins before the addition of Tankyrase 1 (150 μg/mL) and NAD+ (120 μM) followed by further incubation for 30 mins. 3) TRF1 (200 nM) and TIN2L (200 nM) were incubated together for 20 mins without Tankyrase 1 and NAD+. For all three experiments, the protein solutions were mixed with the 200 nM QD-labeled ^BT^pT270 DNA and diluted 50X using the imaging buffer before being introduced into the flow chamber, and further incubated in the flow chamber for 5 mins before video recording. After the third experiment and flushing of the flow chamber, Tankyrase 1 (150 μg/ml) and NAD+ (120 μM NAD+) was introduced into the flow chamber and incubated for 30 mins with the DNA tightropes to monitor the disassembly of the QD-labeled ^BT^pT270 DNA from DNA tightropes.

Protein-QD complexes were excited at 488 nm by a solid-state laser (Sapphire DPSS). The QD signal was split into two channels by a dichroic mirror (T605LPXR, Chroma), and the red signals passed through an optical filter (ET655/40nm, Chroma) before being detected by an EMCCD camera (iXon DU897, Andor Technology). All videos were taken using an inverted microscope (Nikon Ti-E) with a 100X objective (APO TIRF, Nikon) at a time resolution of 50 ms/frame.

### Prism-type total internal reflection fluorescence microscopy (TIRFM) imaging

PEG/biotin-PEG passivated quartz slides were used to construct home-built flow cells, as described previously (68). To avoid nonspecific binding of proteins and DNA inside the imaging chamber, we conducted PEGylation of the cover slide twice (69), and incubated the chamber with 1X Blocking Reagent (Sigma, 11096176001) and 5% Tween-20 (Sigma, P7949) for 10 min (70). The slides were functionalized by incubating the imaging chamber with 0.1 mg/mL streptavidin. Cy5- and biotin-labeled duplexed telomeric DNA containing 6 TTAGGG repeats (69 bp) was then attached to the slide surface through streptavidin-biotin interactions. After incubation of proteins with surface anchored telomeric Cy5-DNA, 5 nM telomeric Cy3-DNA or control DNA without biotin (59 bp) was mixed with the indicated amount of proteins and injected into the imaging chamber. The samples were imaged using a prism-type TIRF microscope. Cy3 and Cy5 excitations were achieved using 532 nm and 640 nm lasers, respectively. The emission from fluorophores was collected through a water immersion objective (60X, 1.2 N.A.), and the signal was split by a Dual-View optical splitter with a 645 nm dichroic mirror. The green and red signals then passed through optical filters (585/70 bandpass filter for Cy3, 655 long-pass filter for Cy5) before being detected by an EMCCD camera (Cascade 512B, Photometrics). Movies at 100 ms/frame were collected using the following excitation sequence: 1) Brief excitation of the Cy5 fluorophore (~ 2 s) to locate DNA molecules; 2) Excitation of the Cy3 fluorophore (~2 min) to monitor DNA molecules bridged by proteins; 3) Brief excitation of the Cy5 fluorophore (~2 s) to reveal whether Cy5 was photobleached. All experiments were performed at room temperature in an imaging buffer containing 20 mM HEPES, pH7.9, 100 mM NaCl, 0.3 mM MgCl_2_, 1 mM EDTA, 0.5 mM DTT, and 2% glucose (w/v), with the addition of an oxygen scavenging system containing100 U/ml glucose oxidase, 1000 U/ml catalase and the triplet-state quenching reagent (2 mM Trolox) (71). Each data set containing more than 2000 traces was collected from at least three independent experiments. Reported values are the average of the independent repeats, and error bars report the SEM of the independent repeats.

### Statistical Analysis

Data obtained from AFM, TIRFM imaging, and the DNA tightrope assay were from at least two to three independent experiments. The number reported are mean ± SEM, unless stated otherwise. The statistical significance level based on the Student’s t-test was set at p<0.05.

## ACKNOWLEDGEMENTS

We would like to thank the Riehn and Xu groups at North Carolina State University, the Opresko group at the University of Pittsburgh, and the Smith group at New York University for technical support.

## AUTHOR CONTRIBUTIONS

H. P. designed and carried out the experiments, analyzed the data, and wrote the paper. P. K., M. L., C. M., R. B., Q. T., P. H., and C. Y. carried out experiments and analyzed the data. D. B. analyzed the data. J. P. designed the experiments. K. W., R. R., S.S., P. L. P, and H. W. designed the experiments and wrote the paper.

