## Supplemental Figure S1 to S11 for "TIN2 facilitates TRF1-mediated *trans*- and *cis*-interactions on physiologically relevant long telomeric DNA"

### Table of contents

**Figure S1. SDS-PAGE of TRF1, TIN2S, and TIN2L purified from insect cells and quantification of their oligomeric states using AFM volume analysis.**

**Figure S2. TIN2S and TIN2L interact with TRF1 on telomeric DNA.**

**Figure S3. TRF1-TIN2 complexes bind specifically to the telomeric region on linear T270 fragments.**

**Figure S4. AFM volume analysis of TRF1-TIN2 bridging multiple fragments of T270 DNA.**

**Figure S5. Control experiments using TIRFM imaging to evaluate Cy5-Cy3 DNA bridging efficiency by TRF1-TIN2 complexes.**

**Figure S6. Lifetimes of transient DNA-DNA pairing events mediated by TRF1 alone.**

**Figure S7. Dual-color differentially labeled TRF1 (green QDs) and TIN2S (red QDs) on T270 tightropes.**

**Figure S8. Quantification of QD labeling efficiency on $^{BT}$pT270 and $^{BT}$noTel DNA.**

**Figure S9. Colocalization of QD-labeled TIN2S and $^{BT}$pT270 DNA on T270 tightropes.**

**Figure S10. Tankyrase 1 removes TRF1 from the telomeric DNA and TIN2S protects TRF1 from Tankyrase 1.**

**Figure S11. Disruption of TRF1-TIN2-mediated DNA-DNA bridging by Tankyrase 1 depends on the presence of NAD+.**

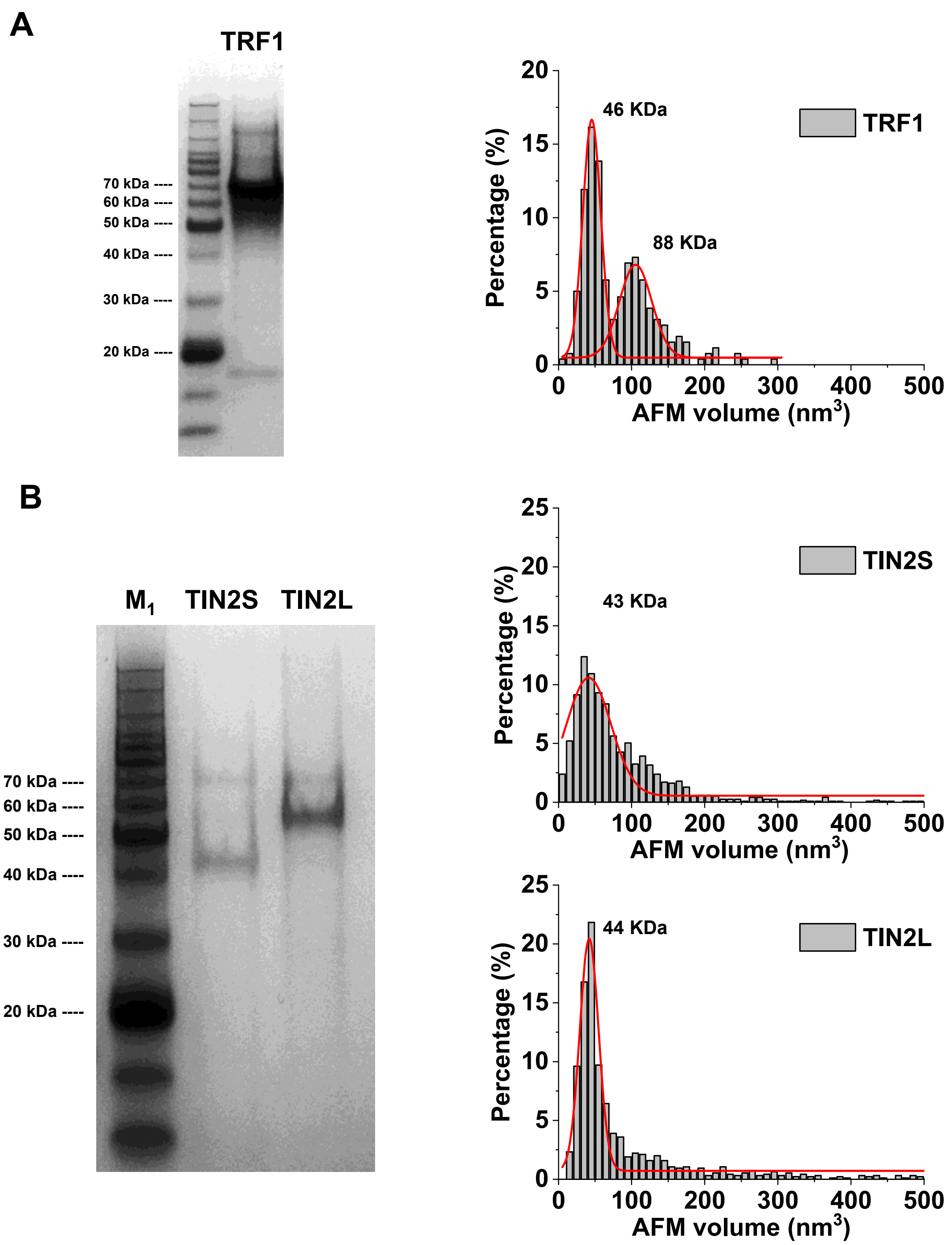


**Figure S1. SDS-PAGE of TRF1, TIN2S, and TIN2L purified from insect cells and quantification of their oligomeric states using AFM volume analysis.**
(**A**) SDS-PAGE of TRF1 (left panel) and AFM volumes of TRF1 (right panel). Fitting the data with a double Gaussian function shows peaks centered at 45.5 ± 12.3 $nm^3$ (mean ± SD) corresponding to monomers (51 kDa), and 105.4 (± 22.2) $nm^3$ corresponding to dimers (102 kDa, $R^2 > 0.98$, N=248). (**B**) SDS-PAGE of TIN2 (left panels) and AFM volumes of TIN2 (right panel). $M_{1:}$ Protein marker (GenScript). HA-TIN2S (39.4 kDa, 2 μg) and TIN2L (50.0 kDa, 2 μg). The red lines in the right panels are Gaussian fits to the data with peaks centered at 41.3 $nm^3$ (±28.3 $nm^3$) for TIN2S ($R^2 > 0.89$, N=1240), and 41.9 $nm^3$ (±12.8 $nm^3$) for TIN2L ($R^2 > 0.94$, N=948). The numbers next to the peaks are the predicted molecular weights based on the equation relating AFM volumes (V) and molecular weights (MW): V ($nm^3$) = 1.45 MW – 21.59 (Kaur *et al.*, Scientific Reports, 2016).

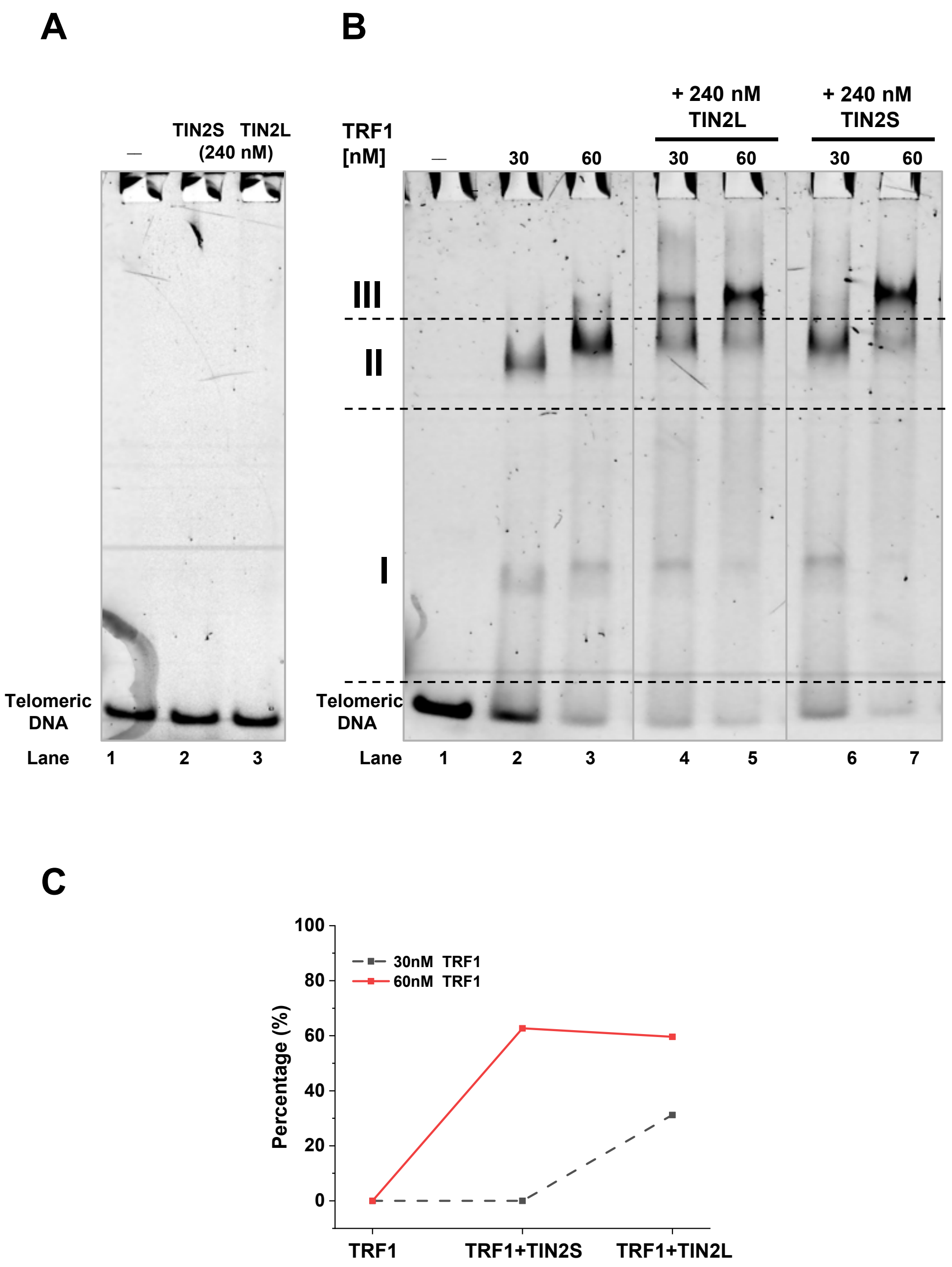


**Figure S2. TIN2S and TIN2L interact with TRF1 on telomeric DNA.**
(**A**) TIN2S or TIN2L alone did not directly bind to Alexa 488-labeled dsDNA containing 3 TTAGGG repeats (5 nM). (**B**) Mobility shift induced by the binding of TRF1 or TRF1-TIN2 complexes to Alexa 488-labeled dsDNA containing 3 TTAGGG repeats (5 nM). Lane 1: DNA only; lanes 2-3: DNA with TRF1 ; lanes 4-5: DNA with TRF1 and TIN2L (240 nM); lanes 6-7: DNA with TRF1 and TIN2S (240 nM). Complexes were categorized as free DNA, complex I, complex II and complex III. The pictures were from the same gel. (**C**) Quantification of the percentage of the complex III formation in the presence of TRF1 alone (30 nM or 60 nM) or TRF1-TIN2 at different TRF1 concentrations (TIN2:240 nM; TRF1: 30 nM or 60 nM).

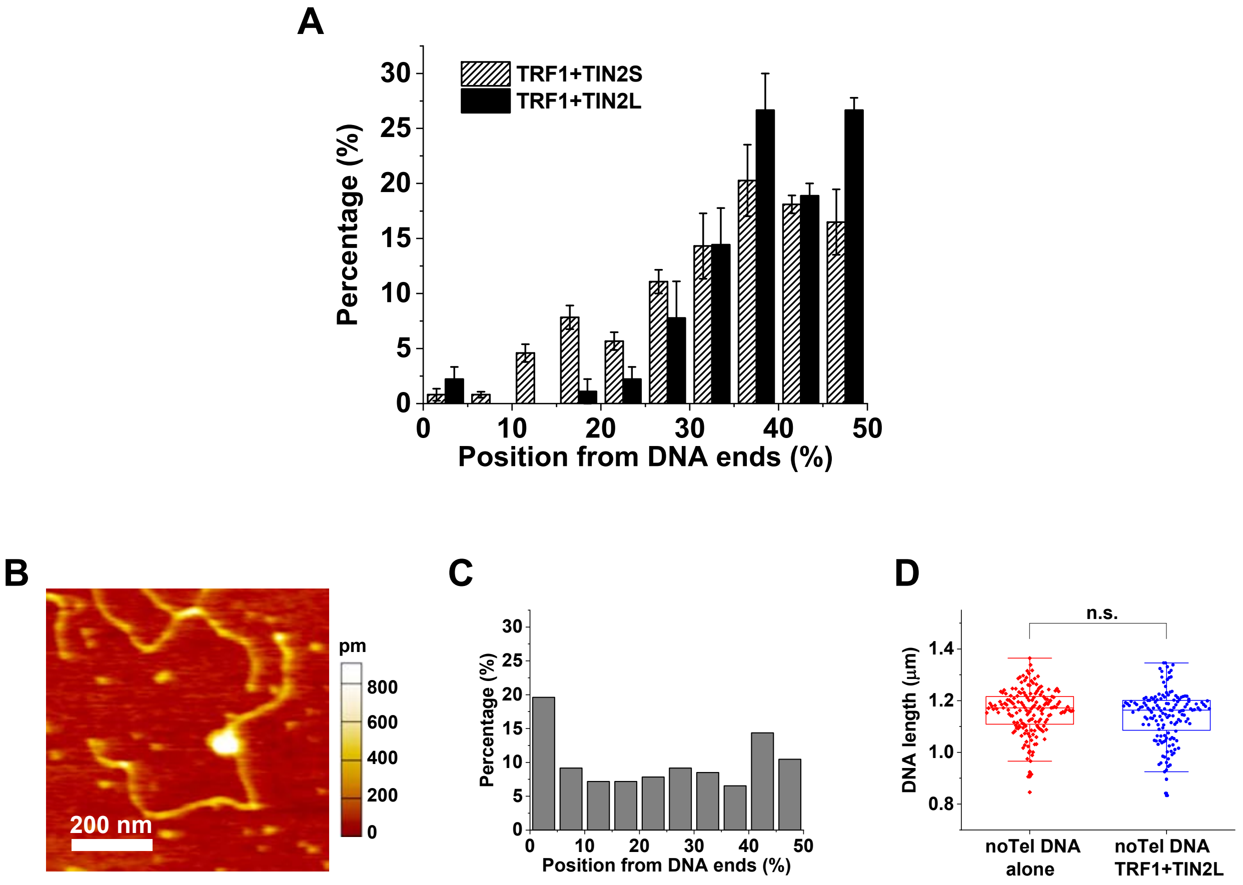


**Figure S3. TRF1-TIN2 complexes bind specifically to the telomeric region on linear T270 fragments.**
(**A**) Binding position analysis showing TRF1-TIN2 complexes specifically binding to the telomeric regions on the linear T270 substrate. Most of the protein complexes (75%, N=88 complexes for TRF1-TIN2L, and 55%, N=185 complexes for TRF1-TIN2S) formed at the $(TTAGGG)_{270}$ region (35%-50%) on T270 DNA. (**B**) AFM image of TRF1-TIN2L binding on the linear non-telomeric (noTel) control DNA (4.1 kb). (**C**) Binding position analysis of TRF1-TIN2L on the control noTel DNA showing nonspecific binding along internal sites along the linear DNA (N=153). (**D**) DNA length measurement showing no significant compaction upon TRF1-TIN2L binding to noTel DNA. The length of DNA alone was 1.16 (± 0.09) μm (N=173), while the length for DNA with TRF1-TIN2L was 1.14 (± 0.10) μm (N=152).

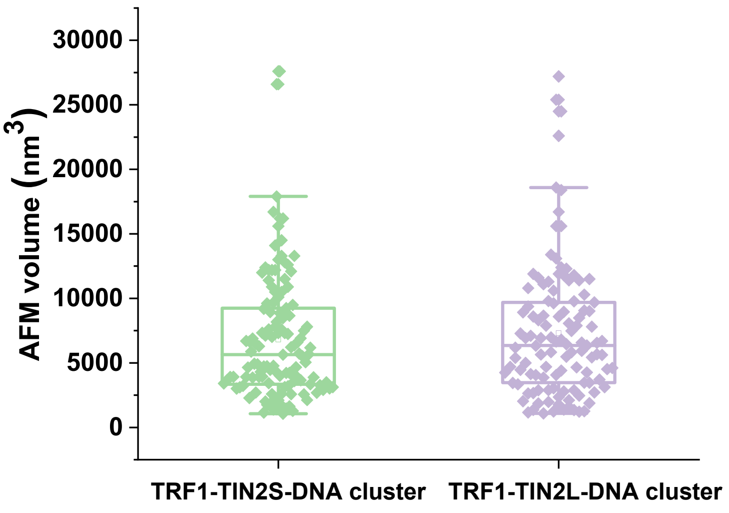


**Figure S4. AFM volume analysis of TRF1-TIN2 bridging multiple fragments of T270 DNA.**

TRF1-TIN2-DNA complexes that bridged multiple fragments of linear T270 DNA (clusters) exhibited board AFM volume distributions. AFM volumes for TRF1-TIN2S-DNA clusters: 6816 ± 4788 nm$^3$ (N=123); AFM volumes for TRF1-TIN2L-DNA clusters: 7286 ± 5229 nm$^3$ (N=109). Examples of TRF1-TIN2-DNA clusters are shown in **Figure 2**.

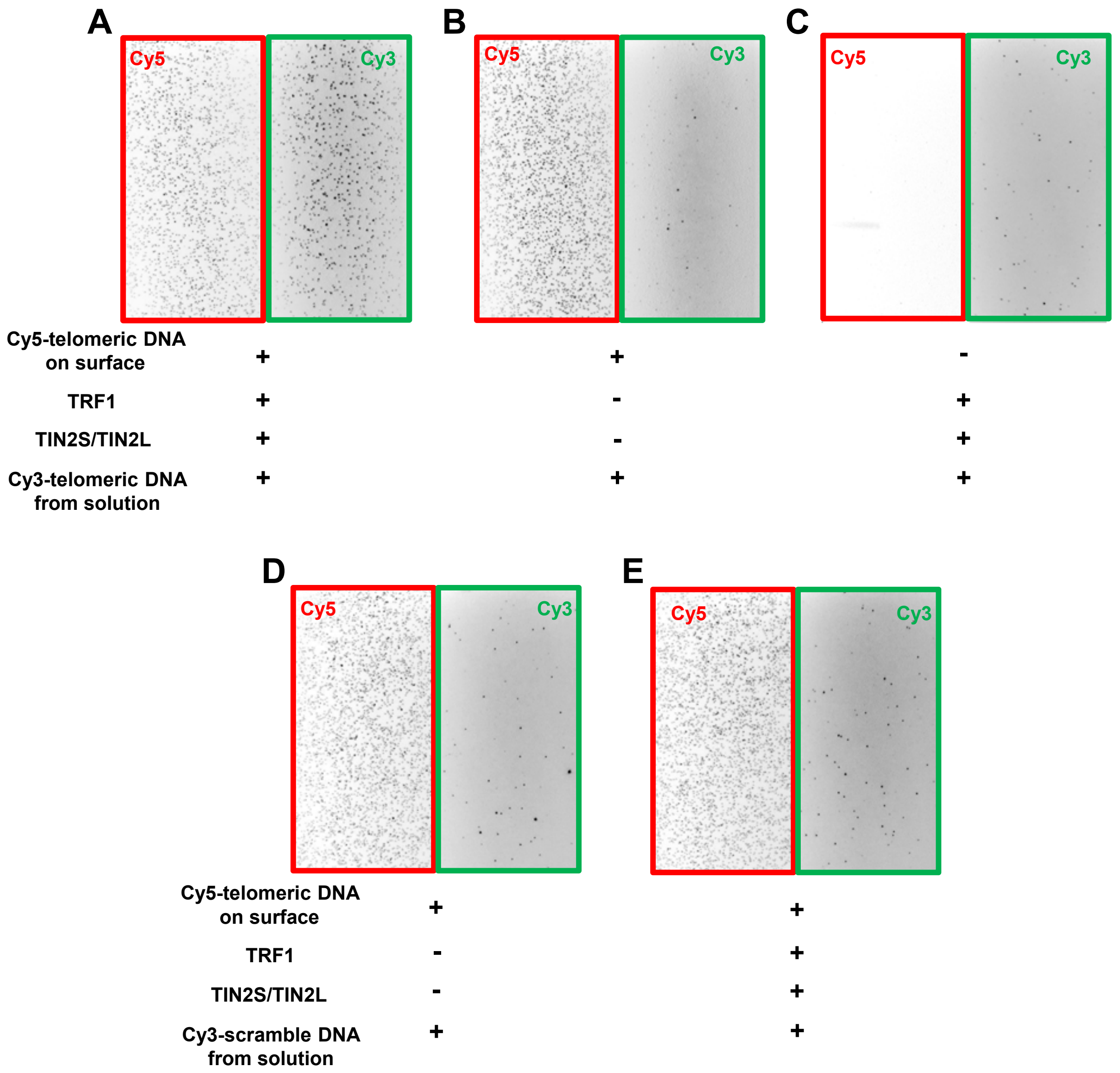


**Figure S5. Control experiments using TIRFM imaging to evaluate Cy5-Cy3 DNA-DNA bridging specificity by TRF1-TIN2 complexes.**
(**A-E**) Representative images from the Cy5 (left) and Cy3 (right) channels under different incubation conditions. (**A** and **B**) With the telomeric Cy5-DNA anchored on the surface and the telomeric Cy3-DNA in solution. (**A**) With the addition of both TRF1 and TIN2 in the imaging chamber. (**B**) Without TRF1-TIN2 in the imaging chamber. (**C**) Without the surface anchored telomeric Cy5-DNA, but with the telomeric Cy3-DNA and proteins (TRF1 and TIN2) in the imaging chamber. (**D** and **E**) With surface anchored telomeric Cy5-DNA and control Cy3-DNA with scrambled sequences in solution. (**D**) Without proteins. (**E**) With TRF1 and TIN2. For all control experiments, Cy3-Cy5 DNA colocalization signals were less than 5% of total traces.

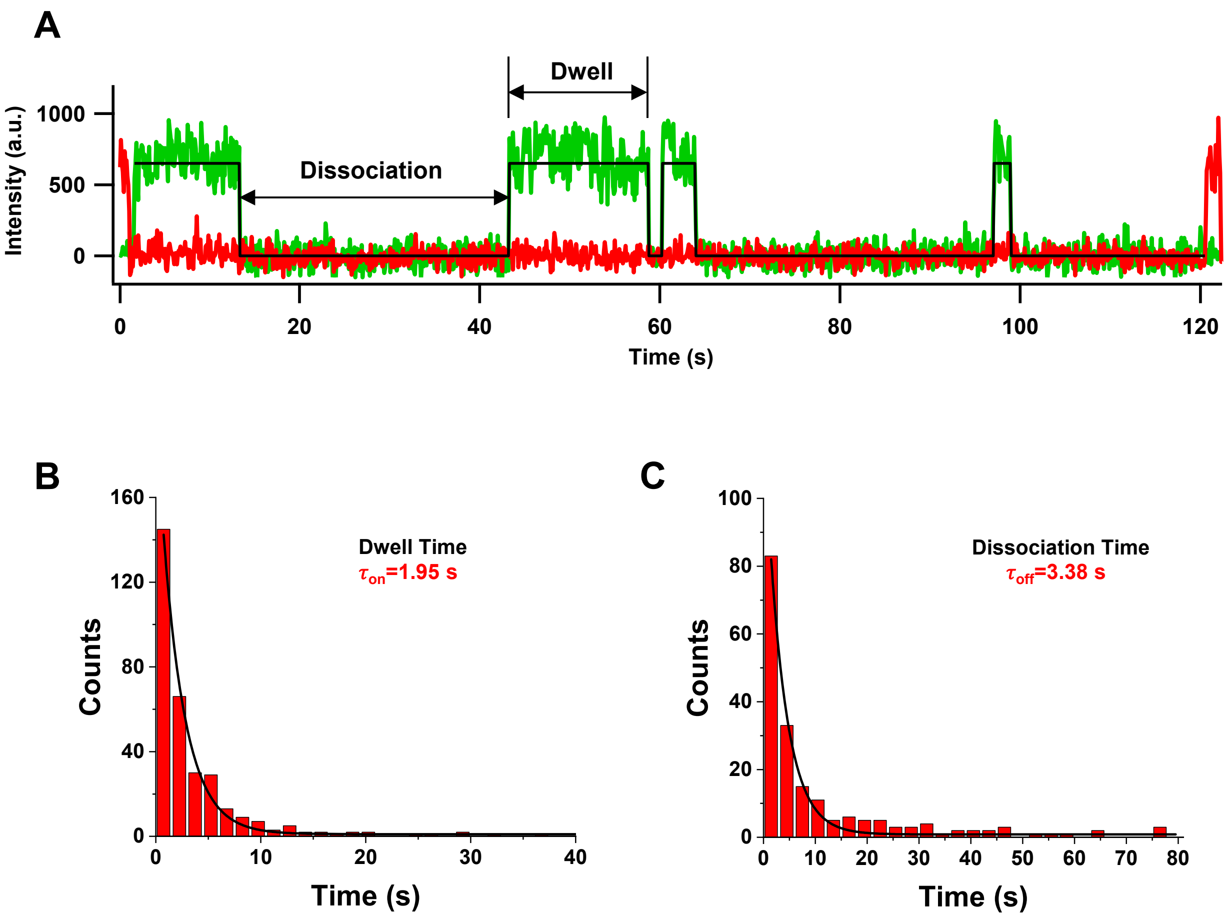


**Figure S6. Lifetimes of transient DNA-DNA pairing events mediated by TRF1 alone.**

(**A**) A representative trace showing telomeric Cy5-Cy3 DNA-DNA pairing events induced by TRF1. The black line is fitting by the Chung-Kennedy model to identify the “on” and “off” state. (**B** and **C**) Dwell time (**B**) and dissociation time (**C**) distributions for TRF1-mediated telomeric Cy3-Cy5 DNA bridging events. Fitting with single exponential functions ($R^2$>0.98) provides the dwell time ($\tau_{on}$=1.95 s, N=322 ) and dissociation time ($\tau_{off}$=3.38 s, N=191).

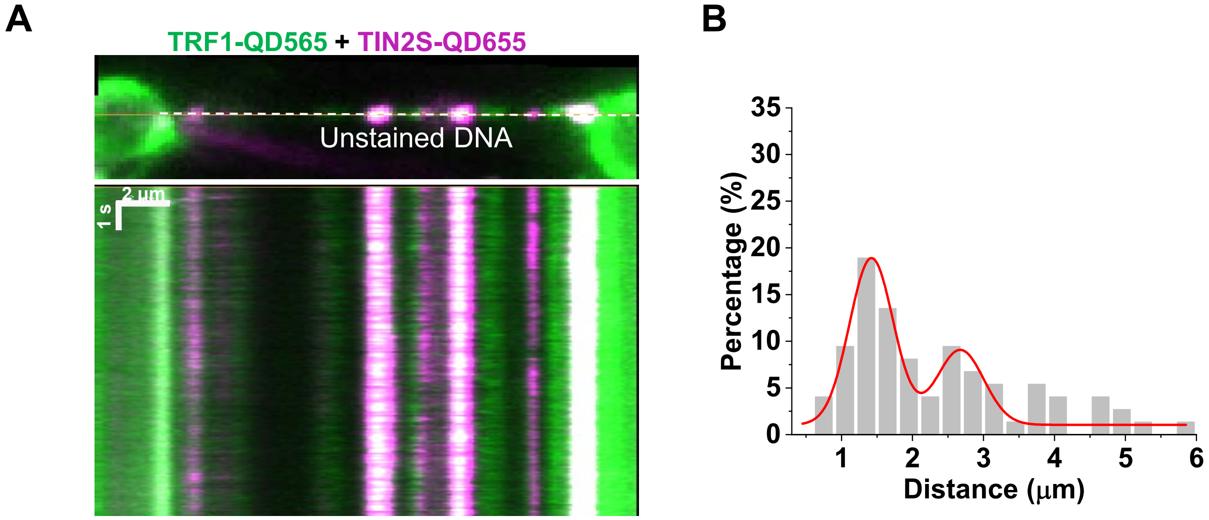


**Figure S7. Dual-color differentially labeled TRF1 (green QDs) and TIN2S (red QDs) on T270 tightropes.**
There were 63.5% green (565 nm) QD-labeled molecules (TRF1, N=215), 0.5% red (655 nm) QD labeled (TIN2S, N=2), and 36.0% dual color-labeled complexes (TRF1+TIN2S, N=122). Right panel: the distance between two adjacent TIN2S-QDs on T270 DNA tightropes. Fitting the data with a double Gaussian function (solid red line) shows peaks centered at 1.42 μm and 2.67 μm (N=79, $R^2$> 0.87).

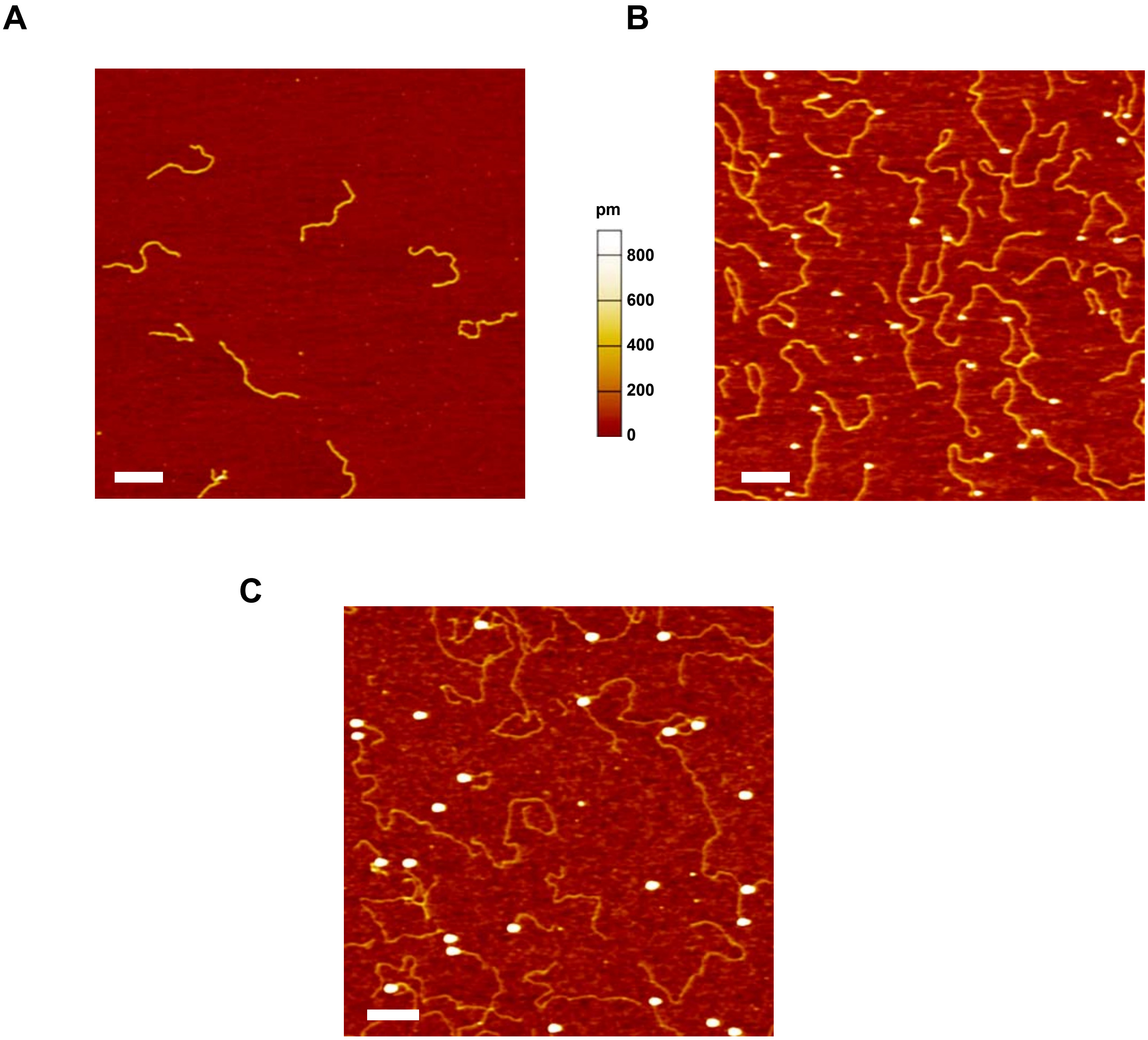


**Figure S8. Quantification of QD labeling efficiency on $^{BT}$pT270 and $^{BT}$noTel DNA.**
(**A** and **B**) AFM images of $^{BT}$pT270 DNA in the absence (**A**) and presence of strep-QDs (**B**). 60.7% (± 3.1%) of $^{BT}$pT270 DNA molecules were labeled with red (655 nm) strep-QDs (N=354). (**C**) AFM image of $^{BT}$noTel DNA (4.1 kb) in the presence of strep-QDs. 67.7% (± 6.7%) of $^{BT}$noTel DNA molecules were labeled with red (655 nm) strep-QDs (N=308). The ratio of DNA and QDs for labeling was 1:10 (400 nM: 4 μM). XY scale bars: 200 nm.

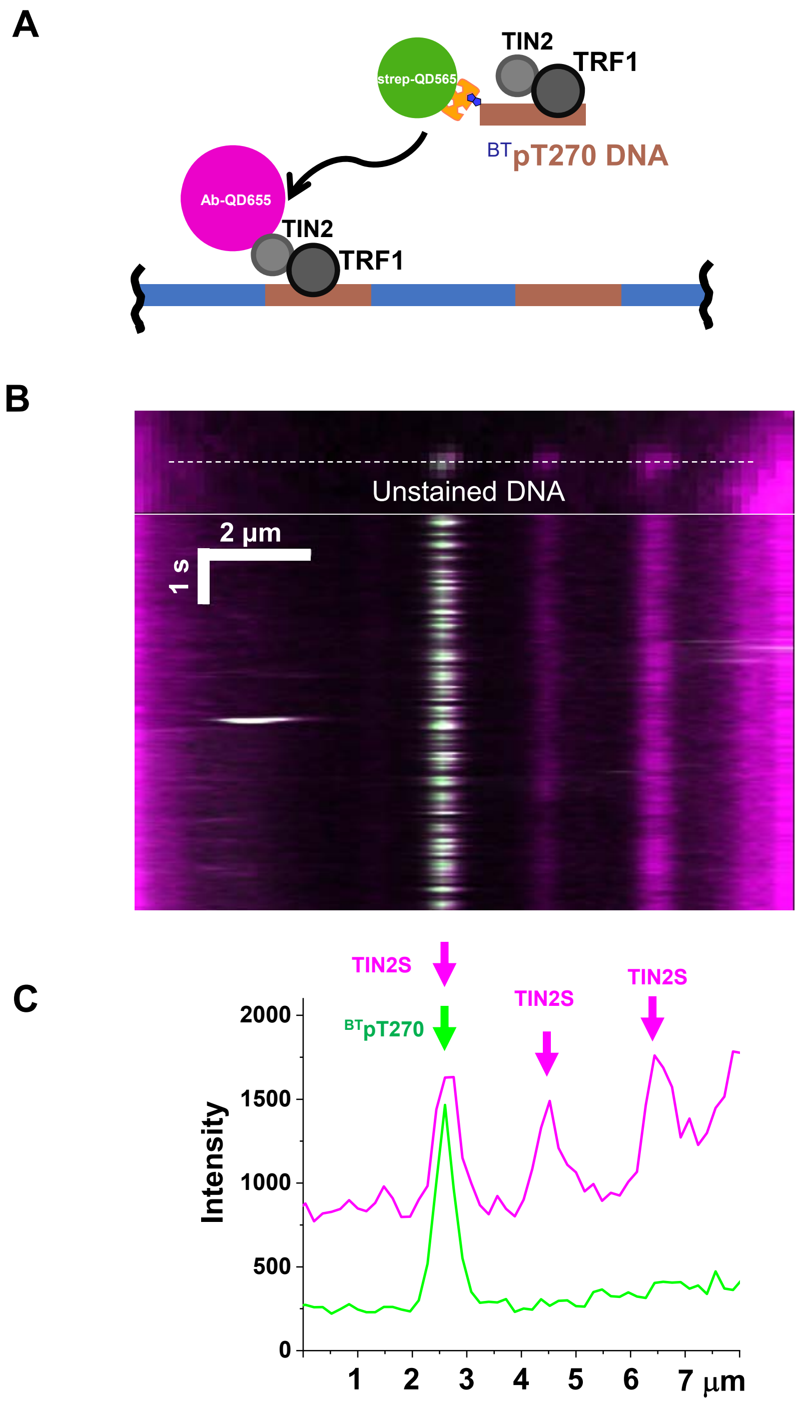


**Figure S9. Colocalization of QD-labeled TIN2S and $^{BT}$pT270 DNA on T270 tightropes.**
(**A**) Schematics of the bridging of the $^{BT}$pT270-QD and T270 tightropes by TRF1-TIN2S. TIN2S was labeled with HA-Ab-QDs. (**B**) Representative fluorescence image and kymograph showing colocalization of red (655 nm) QD-labeled TIN2S and green (565 nm) QD-labeled $^{BT}$pT270 on the T270 tightrope. (**C**) Fluorescence intensity profile showing the colocalization of TIN2S and $^{BT}$pT270. The majority of $^{BT}$pT270 (~87%, N=28) bridged to T270 DNA tightropes colocalized with TIN2-QDs.

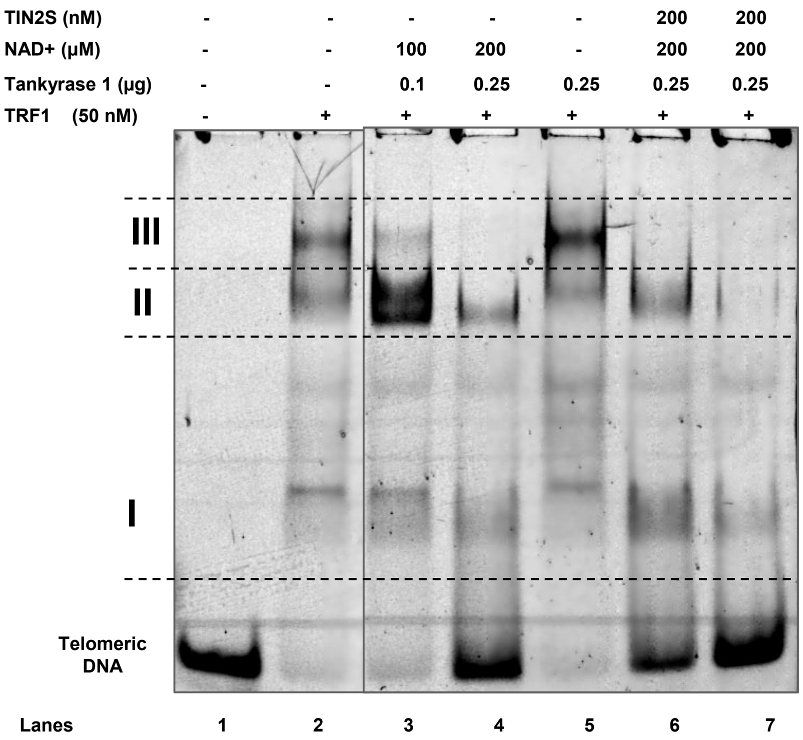


**Figure S10. Tankyrase 1 removes TRF1 from the telomeric DNA and TIN2S protects TRF1 from Tankyrase 1**. EMSA of telomeric DNA in the presence of TRF1 only (lane 2), TRF1 with increasing concentrations of Tankyrase 1 and NAD+ (lanes 3 and 4), the negative control without NAD+ (lane 5), TRF1+TIN2S+Tankyrase 1+NAD+ (lanes 6 and 7). Lane 6: TIN2S was incubated with TRF1 first for 25 mins before the addition of Tankyrase 1. Lane 7: TRF1 was incubated with Tankyrase 1 first for 25 mins before the addition of TIN2S. All the lanes were compiled from the same gel.

**TRF1 + TIN2L + Tankyrase 1 + $^{BT}$pT270**

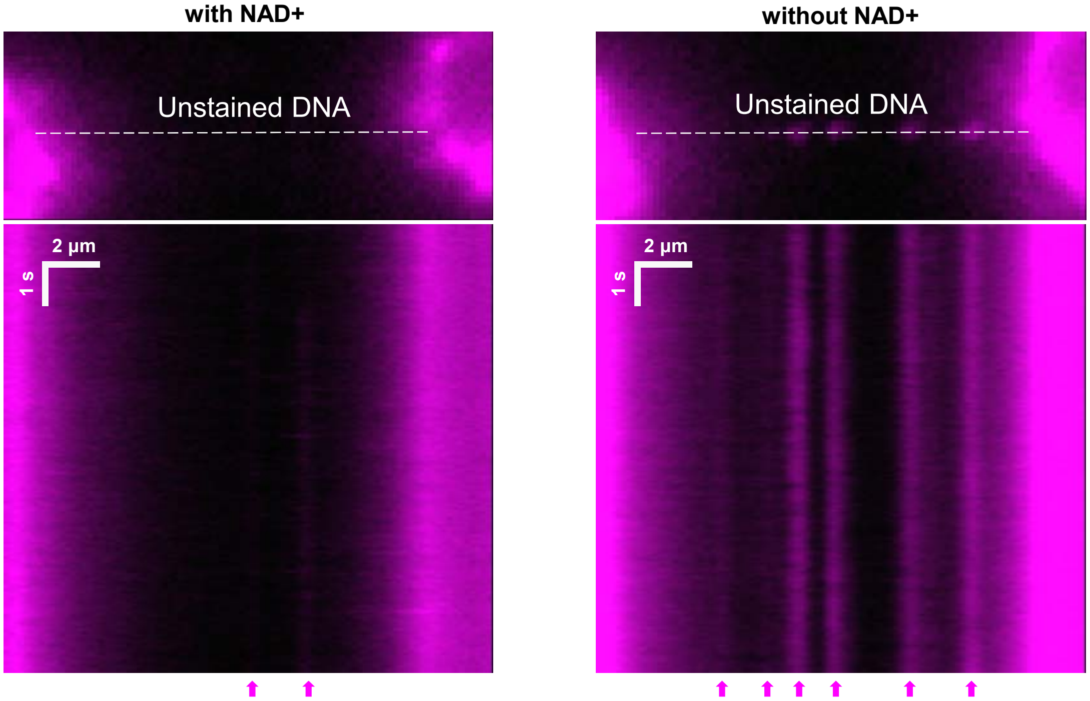


**Figure S11. Disruption of TRF1-TIN2-mediated DNA-DNA bridging by Tankyrase 1 depends on the presence of NAD+.**
Videos were taken over a same area containing a DNA tightrope under two experimental conditions either with or without NAD+. Left: TRF1 + Tankyrase 1 + TIN2L + $^{BT}$pT270 DNA fragments labeled with QDs, and NAD+ (120 μM), total count 20 events from 10 videos; Right: TRF1 + Tankyrase 1 + TIN2L + $^{BT}$pT270 DNA fragments labeled with QDs, but without NAD+, total count 203 events from 10 videos.
